# Ultra-high resolution fMRI reveals origins of feedforward and feedback activity within laminae of human ocular dominance columns

**DOI:** 10.1101/2020.05.19.102186

**Authors:** Gilles de Hollander, Wietske van der Zwaag, Chencan Qian, Peng Zhang, Tomas Knapen

## Abstract

Ultra-high field MRI can functionally image the cerebral cortex of human subjects at the submillimeter scale of cortical columns and laminae. Here, we investigate both in concert, by, for the first time, imaging ocular dominance columns (ODCs) in primary visual cortex (V1) across different cortical depths. We ensured that putative ODC patterns in V1 (a) are stable across runs, sessions, and scanners located in different continents (b) have a width (∼1.3 mm) expected from post-mortem and animal work and (c) are absent at the retinotopic location of the blind spot. We then dissociated the effects of bottom-up thalamo-cortical input and attentional feedback processes on activity in V1 across cortical depth. Importantly, the separation of bottom-up information flows into ODCs allowed us to validly compare attentional conditions while keeping the stimulus identical throughout the experiment. We find that, when correcting for draining vein effects and using both model-based and model-free approaches, the effect of monocular stimulation is largest at deep and middle cortical depths. Conversely, spatial attention influences BOLD activity exclusively near the pial surface. Our findings show that simultaneous interrogation of columnar and laminar dimensions of the cortical fold can dissociate thalamocortical inputs from top-down processing, and allow the investigation of their interactions without any stimulus manipulation.

**Significance Statement:** The advent of ultra-high field fMRI allows for the study of the human brain non-invasively at submillimeter resolution, bringing the scale of cortical columns and laminae into focus. De Hollander et al imaged the ocular dominance columns and laminae of V1 in concert, while manipulating top-down attention. This allowed them to separate feedforward from feedback processes in the brain itself, without resorting to the manipulation of incoming information. Their results show how feedforward and feedback processes interact in the primary visual cortex, highlighting the different computational roles separate laminae play.

## Introduction

Ultra-high field functional MRI (UHF-fMRI, at field strengths of 7 Tesla or more) is transforming the field of human neuroimaging (Marques and Norris, 2018; Trattnig et al., 2018; van der Zwaag et al., 2016). By increasing signal-to-noise ratio (SNR) and spatial resolution, UHF-fMRI allows researchers to non-invasively study cortical functional organization at the mesoscopic scale of cortical columns and layers in healthy humans (De Martino et al., 2018; Dumoulin et al., 2018). Perhaps the most well-known form of cortical organization at the mesoscopic scale is columnar: ocular dominance columns (ODCs) are patches of cortex that are predominantly sensitive to input from only one of the two eyes, forming columns perpendicular to the surface of primary visual cortex, V1 (; Hubel and Wiesel, 1969; Tootell et al., 1988; Adams et al., 2007; Adams and Horton, 2009; Dougherty et al., 2019). In addition, models of cortical microcircuits contend that the organization orthogonal to that of cortical columns, along cortical depth, separates input-driven and feedback activity in primary sensory regions (Bastos et al., 2012; De Martino et al., 2018; Kuehn and Sereno, 2018; Rockland and Pandya, 1979; Self et al., 2019; Stephan et al., 2019). Specifically, thalamocortical “feedforward” inputs are believed to mostly reside at the “middle”, granular layer (Bastos et al., 2012; Stephan et al., 2019) and, “deep”, infragranular layers (Constantinople and Bruno, 2013). Feedback processes, like perceptual predictions and attention, however, are believed to mostly signal into supragranular and infragranular layers (Rockland and Pandya, 1979; Self et al., 2019). Therefore, combined laminar-columnar UHF-fMRI could dissociate BOLD activity in V1 due to thalamocortical *input* (e.g., by monocular stimulation of the preferred eye of a given ODC) from activity due to *higher order processes* (e.g, by manipulating attention) in the same cortical columns (Schneider et al., 2019). Dissociating feed-forward and feedback processes that deal with ocularity information provides a potential opportunity to empirically validate the wiring of cortical microcircuit models non-invasively in healthy human subjects, and elucidate the underlying mechanisms of a range of phenomena such as binocular rivalry (Brascamp et al., 2018; Wheatstone, 1838) and stereoscopic depth vision (Goncalves and Welchman, 2017).

Earlier studies that aimed to show the laminar profile of *feedforward* thalamocortical inputs using UHF-fMRI did so by manipulating stimulus features, such as the presence of a stimulus (Koopmans et al., 2010) or stimulus contrast (Lawrence et al., 2019). Arguably, such stimulus manipulations could still induce higher-order feedback processes, in addition to feedforward input, because the appearance of such stimuli are inherently different and can thereby change feedback processes into V1 as well (Petro et al., 2014). Here, we leverage the preferred ocularity of mapped ODCs to infer whether monocular stimulation in the left or right eye is increasing thalamocortical input for a given patch of cortex – without the need to resort to stimulus manipulations.

Concretely, we presented visual stimuli while fully controlling which eye received visual stimulation, allowing us to contrast left-eye and right-eye responses and map ocular preference across V1. We then independently manipulated the relevance of monocular or binocular information for our participants: participants were asked to report on changes in either a binocularly presented fixation mark, or a monocularly presented rotating-checkerboard stimulus. Importantly, the objective visual stimulation was identical across these two attentional conditions. We used 7T gradient-echo (GRE-)fMRI and isotropic voxels at a very high resolution (Fig 2A, 0.7 mm isotropic; 0.343 mm^3^) to sample activation patterns within the cortical ribbon at different cortical depths (Bazin et al., 2014; Dale et al., 1999; Fischl et al., 1999; Polimeni et al., 2018). This allowed us to quantify ODC patterns across cortical depth, and investigate how these patterns change as a function of thalamocortical input and stimulus relevance (attentional state). By using isotropic voxels, we could also reduce the number of voxels that overlap with draining vasculature near the pial surface, thereby increasing the effective resolution on the cortical surface (Kay et al., 2019). Finally, combining ODC measurements with population receptive field estimates at all locations allowed us to relate the ocular dominance column patterns to their place in the retinotopic visual field (Adams and Horton, 2009; Dumoulin and Wandell, 2008).

**Figure 1:**
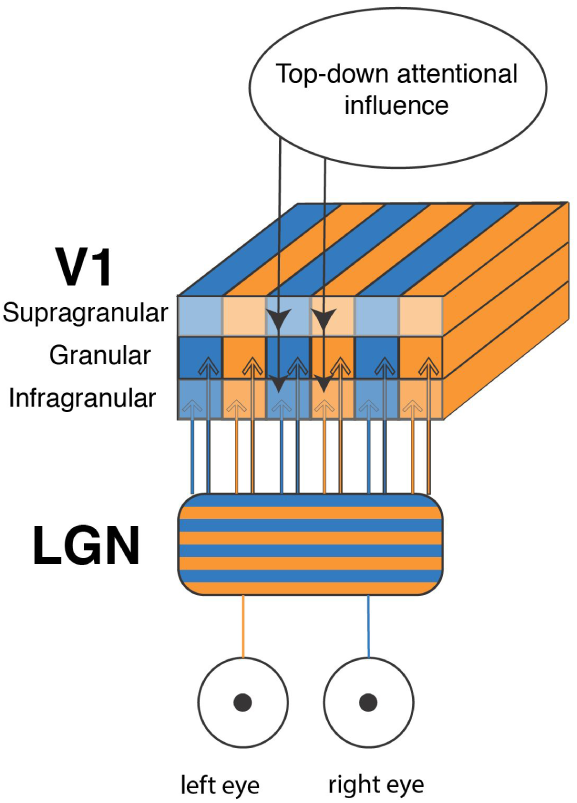
Schematic illustration of laminae and columns in V1. Visual information enters the visual system via the cornea of the left and right eye (bottom), and then, through the optical nerve reaches primary visual (V1) via the lateral geniculate nucleus (LGN; middle) in the thalamus. The signal then predominantly projects into the granular, as well as the infragranular layer in V1. The ocularity of the signal is preserved along so-called “ocular dominance columns” of approximately 1 mm wide. Feedback processes from higher-order cortical regions project largely into supragranular and infragranular layers.

**Figure 2:**
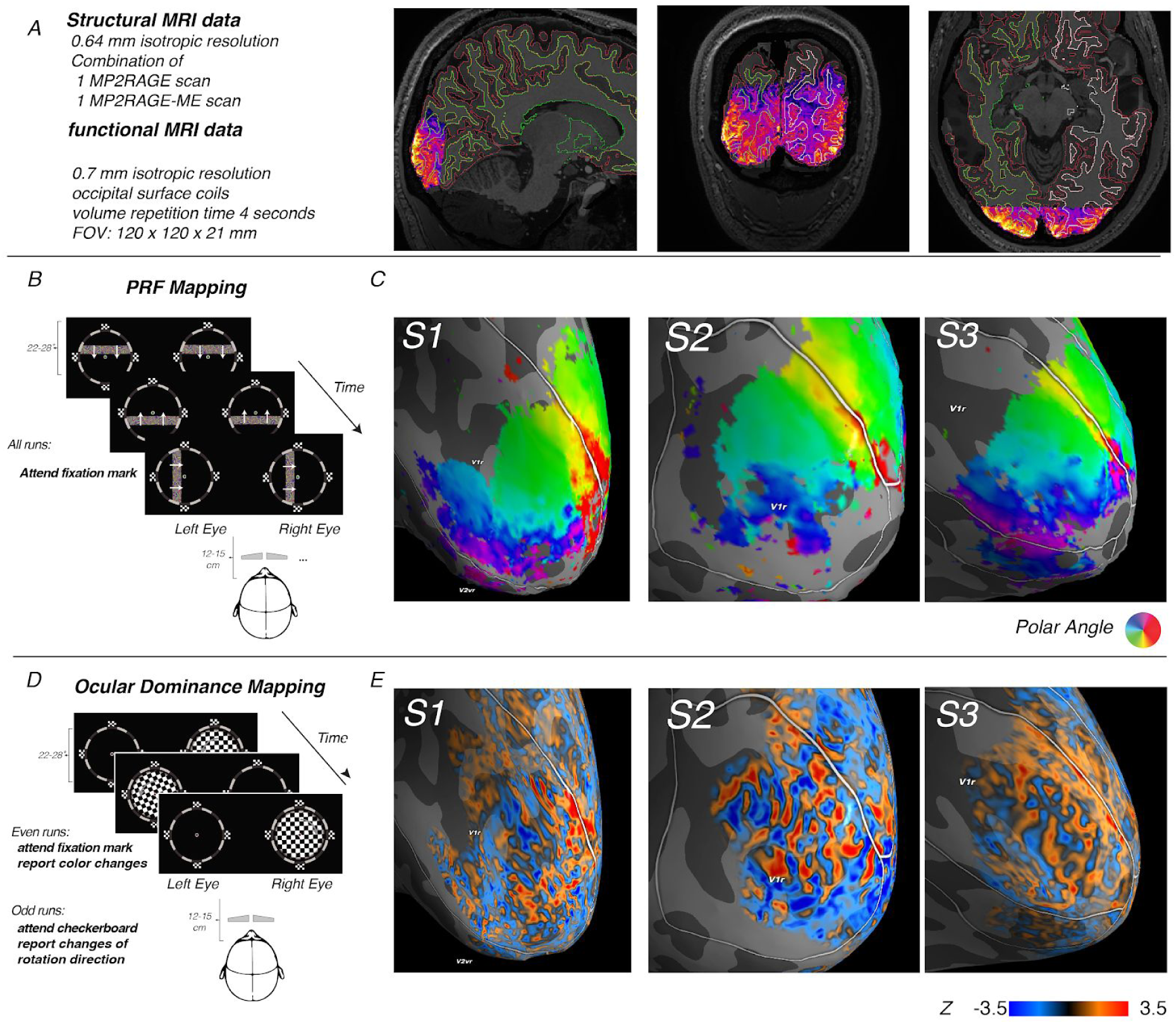
PRF and ocularity maps in V1. ***A) functional MRI data was collected at 7 Tesla with isotropic resolution of 0.7 mm*** and occipital surface coils with a restricted field-of-view, but optimized signal from the calcarine sulcus. Overlaid contour profile shows white (green/white lines) and gray matter segmentations (red lines) created by Freesurfer at 0.64 mm resolution, based on the mean T1-weighted UNI image of a MP2RAGE and MP2RAGE-ME-sequence. The purple/orange image is the mean EPI image of a run registered to the anatomical data using boundary-based registration and illustrates the quality of the registration, as well as the field-of-view. ***B) A standard Population Receptive Field (PRF) mapping paradigm was presented*** to the subjects for 3 runs of approx 5 minutes. In this paradigm, a flickering bar consisting of many gabor patches was moved across the central aperture in 4 cardinal and 4 oblique directions. The projection screen was approximately 12-15 centimeters from the subject’s eyes, which led to an ocular field-of-view of approximately 22-28 degrees-of-visual-angle. ***C) The PRF mapping paradigm yielded individual retinotopic field maps in and around the calcarine sulcus.*** These maps were used for a very fine-grained delineation of the V1/V2-border. Here, we show the retinotopic maps of three representative subjects. ***D) An ocular dominance mapping paradigm with both monocular and binocular stimulus elements was presented to each subject*** for 8-10 runs. Before half of the runs, subjects were instructed to report changes in the color of the binocularly-presented fixation dot; before the other half of the runs, subjects were instructed to report the rotating direction of the checkerboard. ***E) The ocular dominance mapping paradigm yielded a contrast map (left > right eye stimulation)*.** Note how the z-values change between large positive and negative values across the cortical surface. The polar angle maps, as well as the ocularity maps can be visually inspected interactively at http://aeneas.labs.vu.nl/odc/

In the following, we first establish the consistency and robustness of ODC patterns and replicate some key characteristics of ODCs. Then, we focus on the effects of (a) monocular stimulation of the preferred eye versus the non-preferred eye of an ODC and (b) the effect of attending a stimulus that is presented monocularly versus a stimulus that is presented binocularly. The central question of this study was how these effects manifest differently across cortical depth. We hypothesized that the effect of monocular stimulation should be most prevalent at the middle cortical depth, where the granular layers reside (Bastos et al., 2012; but see Constantinople and Bruno, 2013; De Martino et al., 2018; Kuehn and Sereno, 2018; Rockland and Pandya, 1979; Self et al., 2019; Stephan et al., 2019), whereas the effect of attention should be most prevalent at outer cortical depth, where the projections originating from higher-order visual areas are located (Bastos et al., 2012; De Martino et al., 2018; Kuehn and Sereno, 2018; Lawrence et al., 2019; Rockland and Pandya, 1979; Self et al., 2019; Stephan et al., 2019).

Finally, we were also interested to see if there was an *interaction* between the effect of monocular stimulation of the preferred eye and attentional state. In other words: is the effect of attention modulated by the stimulation of the preferred versus non-preferred eye? We speculated that humans might be able to suppress the thalamocortical input into V1 representing information that is not relevant to the task at hand. If this would be the case, we would expect the effect of attention to be larger in ODCs where the preferred eye is stimulated as compared to ODCs where the non-preferred eye is stimulated.

In the following we report robust V1 ODC patterns at all cortical depths. Laminar deconvolution and decoding analyses confirm that ocular-dominance information is indeed most robust at middle cortical depth. The behavioral relevance of the monocular stimulus induced a higher BOLD response but only at outer cortical depth, consistent with the standard model that cortico-cortical feedback projections largely reside in the outer layers. Lastly, there was no significant interaction between these two effects, suggesting that, in our task, thalamocortical input is not modulated by attention in an ocular-specific manner.

## Results

### Univariate contrasts on the surface

We first used population receptive field mapping (PRF; Dumoulin and Wandell, 2008) to delineate V1 on the cortical surface of individual subjects. Figure 2C illustrates the retinotopic maps and V1/V2 segmentations of three representative subjects. Figure 2E shows the z-statistics of the contrast ‘left eye stimulation > right eye stimulation’ in the same three subjects. Qualitative inspection suggests there are very small, significant clusters of ocular specificity that alternate between left- and right-sensitivity at a fine spatial scale.

### Reproducibility of ocular dominance maps

In line with earlier work on fMRI of ODCs (Cheng et al., 2001; Yacoub et al., 2007), we first tested whether ocularity maps were reproducible over time. For six subjects, we split the data into the first and second part of the session and estimated activation patterns for these two halves separately. We correlated the activation patterns of these two halves on the surface within the V1 mask that was defined by the retinotopic mapping paradigm. For this analysis, the activity patterns were averaged across cortical depth. Out of six subjects, five subjects showed robust correlations in their activation pattern. These within-session activity pattern correlations ranged from r = 0.41 – 0.80 (median 0.59; see Figure 3A). The sixth subject (subject 5) showed no robust within-session correlation (r = −0.02 for the left and r = 0.02 for the right hemisphere) and was removed from any subsequent analyses. Another subject’s (subject 1) irregular folding of the left hemisphere placed a large part of the calcarine outside the scan protocol (see supplementary materials S3 for details). This hemisphere was left out of all subsequent analyses. For the five subjects that showed robust correlations, the binarized ocular preference of the surface vertices (‘left’ or ‘right’) was consistent across the two session halves for 60 – 79% of the vertices (median 70%).

**Figure 3:**
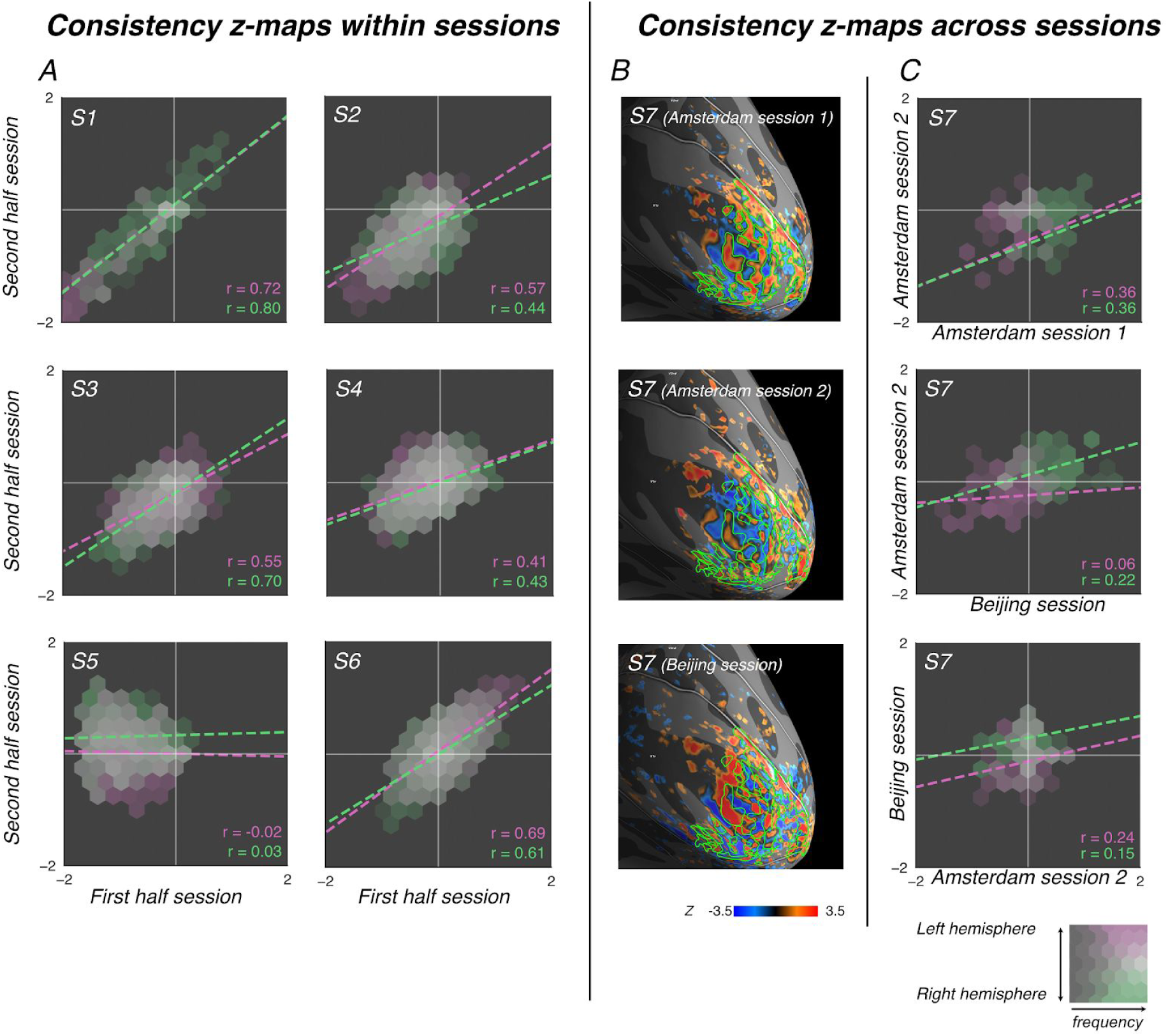
Robustness across runs and sessions. ***A) Correlations between ocularity maps of first and second half of the mapping sessions.*** All subjects except subject 5 show consistent ocularity patterns across the two halves of their session. ***B) One subject was scanned during three separate sessions and showed visually similar ocularity patterns across these three sessions***. One session took place in a different scanner from a different vendor (Siemens) in a different MR center, 7828 kilometers away. ***C) Across both hemispheres and across all three sessions, the ocularity patterns are consistent, albeit less so than within-session ocularity maps***.

For a seventh subject, we acquired data in two separate sessions in Amsterdam, as well as a third session in Beijing, China, using a different 7T MRI system (see methods section for details). Across the two sessions in Amsterdam, activation maps in V1 correlated with r=0.36 in both hemispheres. The activation map of the first session in Amsterdam correlated with the activation map in Beijing with r = 0.06 for the left hemisphere and r=0.22 for the right hemisphere. The activation pattern of the second session in Amsterdam correlated with those from Beijing with r=0.24 for the left hemisphere and r=0.15 for the right hemisphere. Although lower than within-scanner correlations (likely due to residual misregistration and image distortions relating to differences in scanner setups; see Figure 3B-C), these correlations are highly significant (p<0.001).

### ODC properties of ocular-preference maps

Having verified the consistency and robustness of these ocular dominance patterns, we turned to more detailed analyses of the spatial properties of these ocular dominance patterns to positively identify these patterns as ODCs.

First, we know that ODCs are largely constrained to V1, stopping rather abruptly at the V1/V2 boundary (Adams et al., 2007; Adams and Horton, 2009). Indeed, the patterns we find are restricted to the area of V1. The percentage of significant z-values at p < 0.05 (two-sided, absolute z-values higher than 1.96) was higher than chance level in V1, while not in V2 for 5 out of 6 subject-sessions for the left hemisphere (average of 10.7 % vs 5.2 %; t(6) = 2.00, p=0.10) and for 6 out of 7 subject-sessions for right V1 (average of 10.4 % vs 5.3 %; t(7) = 1.55, p=0.17); see Figure 4A).

**Figure 4:**
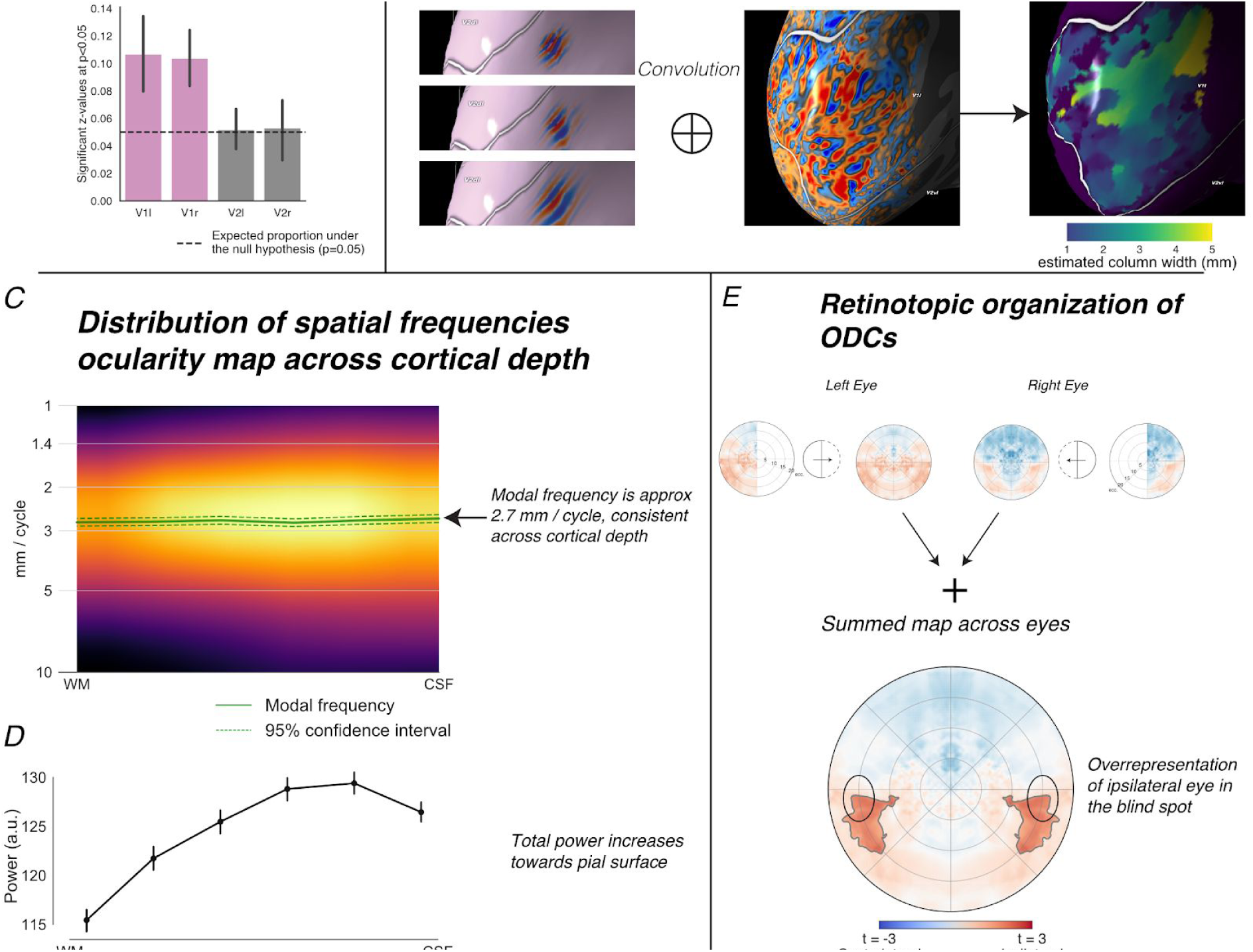
Known properties of ODC that replicate in UHF-fMRI ODC mapping. ***A) In V1, but not V2, the distribution of z-values is robustly different from chance.*** This is consistent with earlier work that has shown that V1 is considerably more sensitive to ocularity than other areas (although there are monocular “stripes” in V2). ***B) Estimation of spatial frequencies on the surface:*** we used a filter bank of Gabor patches and convolved it with ocularity maps on a flattened cortical surface, to estimate a map of main spatial frequencies across V1. ***C) The dominant spatial frequency is constant across cortical depth***. However, other spatial frequencies have more power near the pial surface. ***D) the total amount of power also increases towards the pial surface. E) The individual retinotopic maps of V1 allow for the analysis of the ocularity maps in the visual field.*** Consistent with earlier work, when we combine data from both eyes, we find an overrepresentation of the ipsilateral eye at the approximate visual location of the blind spot (ovals).

Second, ocular dominance columns are known to occur at a specific range of spatial frequencies. Post-mortem work estimates that ODCs are about 730 – 995 micrometers wide (Adams et al., 2007; Adams and Horton, 2009). Yacoub et al. (2007) estimated a column width of 1070 micrometer using GRE and Spin-Echo fMRI techniques and anisotropic voxels. To compare our fMRI results to earlier work, we convolved the left/right-difference z-maps within V1 on a flattened surface with a set of Gabor patches with different spatial frequencies, ranging from 1 mm to 10 mm/cycle, each using 16 different orientations (see Figure 4B). This yielded, for every location on the reconstructed surface, a distribution of power at different spatial frequencies.

All subjects showed a spectral power distribution with a peak around a wavelength of approximately 2.7 mm/cycle (Figure 4C). This is consistent with a column size of approximately 2.7 / 2 = 1.35 mm wide. This is wider than the approximately 1 mm width that is found is post-mortem work. We note that the BOLD transfer function, blurring due to slow k-space sampling and post-hoc resampling to the cortical surface will shift the power spectrum towards longer wavelengths (Chaimow et al., 2018a, 2018b, 2011; Shmuel et al., 2007). The dominant wavelength of 2.7 mm/cycle was constant across cortical depth (F(5, 60) = 0.74, p = 0.60).

Third, as ocular dominance preference is inherited from the bottom-up afferents from the lateral geniculate nucleus (LGN) in the thalamus, the ODC pattern should be strongest in the middle layers of V1 which receive this afferent input. Indeed, the power at the dominant wavelength shows a non-monotonic curve, peaking not at the pial surface (where BOLD amplitude is greatest) but near the middle layers. When we performed Bayesian hierarchical fits of both a linear model and a quadratic model (see methods), the quadratic model explained the relationship between cortical depth and the total spectral power better than the linear model (Watanabe–Akaike information criterion (wAIC) = −398.44 for the quadratic model vs wAIC = −324.53 for the linear model). The peak was estimated to be at approximately 40% cortical depth (95% highest posterior density HPD: [5%, 75%]), where the pial surface corresponds to 0% cortical depth and the gray matter/white matter-border corresponds to 100% cortical depth.

Fourth, we focused on the blind spot: a small part of the visual field of each eye that is occluded by the optic disc (the start point of the optic nerve, where the retina contains no photoreceptors). These blind spots, one for each eye, should be detectable in our ODC patterns as a patch of monocular preference for the ipsilateral eye at about 15 degrees eccentricity, just below the horizontal meridian of the visual field (Adams et al., 2007; Adams and Horton, 2009). To visualize this, we mapped the ODC patterns into visual space using the pRF estimates of our retinotopic mapping experiment (Fig 2B). This indeed revealed areas of dominance of the ipsilateral eye at a location consistent with the blind spot (although somewhat lower in the visual field; peak at x = −/+ 15.1 and y = −7.9 degrees-of-visual-angle; Figure 4E).

We contend that this collection of findings consistently identifies our ocular dominance patterns as human ODCs. We investigated a further set of possible predictions of ODC properties derived from post-mortem studies and animal studies, but technical constraints such as limited spatial resolution and the distortions induced by sampling volumetric data to the surface limit the testability of these predictions (see supplemental materials S2).

### The effect of attention on the BOLD response across cortical depth

After having established that the ocularity maps we found are robust and consistently follow predicted ODC properties, we turned to the question how ODCs in V1 are modulated by spatial attention, across cortical depth. Participants were asked to report on either a monocularly-presented part of the stimulus array (the rotating, flickering checkerboard), or the binocularly-presented part of the stimulus array (the fixation dot) in separate experimental runs featuring identical visual stimulation.

We selected visually responsive voxels in V1 (absolute z-value of *left + right stimulation > baseline* contrast larger than 2.3; approx. 3000 – 9000 voxels per subject) and tested differences in BOLD response amplitude as a function of stimulation and attentional condition. Then, to prevent double dipping, we assigned voxels to be left-eye or right-eye preferring, using leave-two-runs-out cross-validation (one for each attentional condition). Given these ocular preferences, we could now designate voxels as either “stimulated” (e.g., left-eye stimulation of left-preferring column) or “unstimulated” (e.g., right-eye stimulation of left-preferring column). Finally, voxels were assigned to 5 exclusive regions of interest based on cortical depth, as defined by the equivolumetric algorithm of CBS-tools (Bazin et al., 2014; Huntenburg et al., 2018; Waehnert et al., 2014).

Figure 5A shows that, as should be expected when using GRE laminar fMRI (Markuerkiaga et al., 2016; Polimeni et al., 2018; Siero et al., 2011), the BOLD response is larger for more superficial cortical depths, in all conditions (main effect of cortical depth on BOLD response: F(4, 20) = 22.9; p < 0.001 for left V1; F(4, 24) = 17.1; p < 0.001 for right V1).

**Figure 5:**
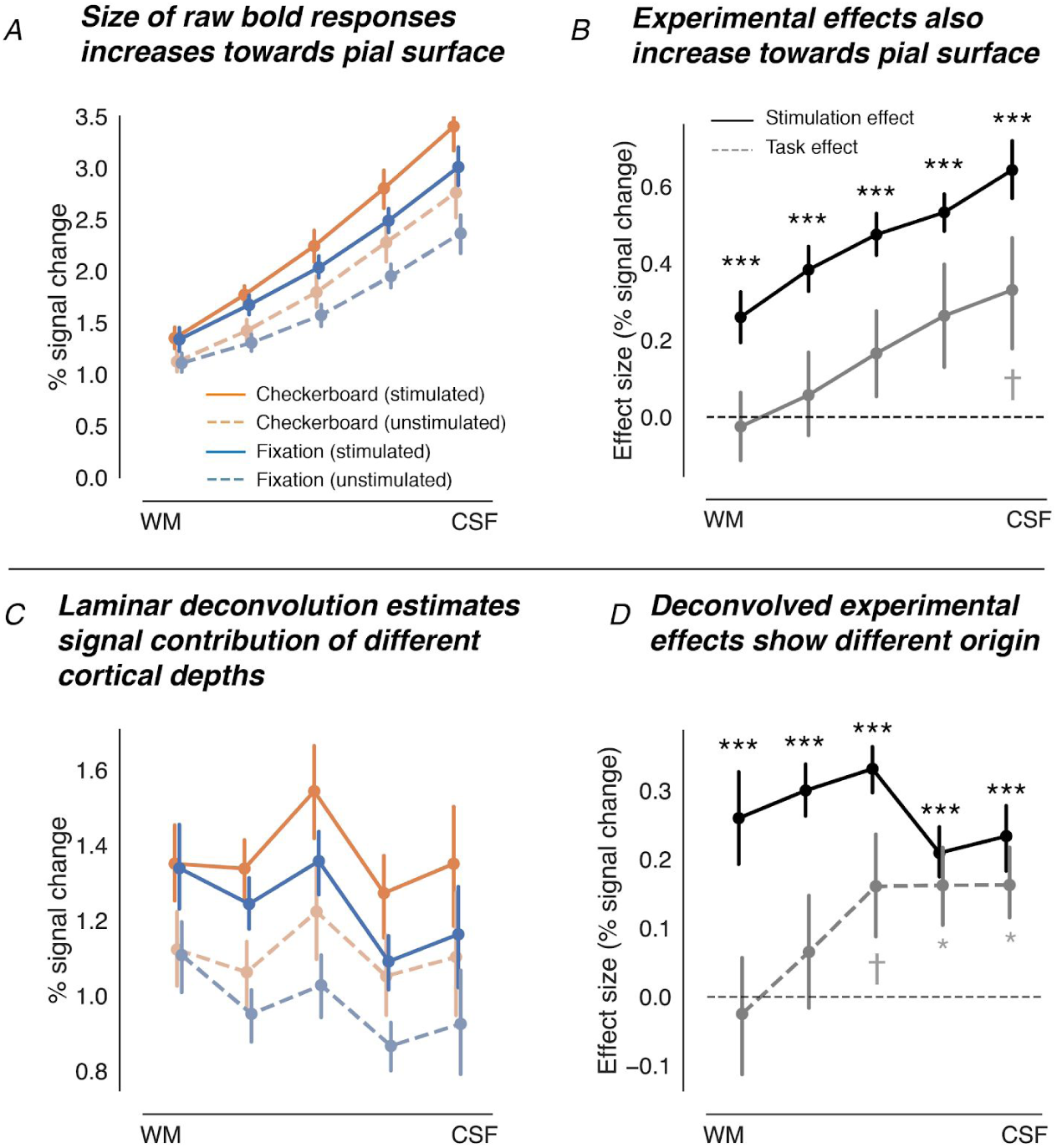
Percent fMRI signal change is modulated by stimulation of preferred eye, attentional condition and cortical depth. **A) *The raw percent signal change increases towards the pial surface*** in all four conditions: with attention on the checkerboard, both stimulated and unstimulated voxels, as well as when the subject is attending the fixation dot for both stimulated and unstimulated voxels. Error bars are standard errors of the mean (SEM), bootstrapped across subjects. **B) *The size of the effects on the attentional and stimulation manipulation also increases towards the pial surface.*** However, stimulating the preferred eye led to increased BOLD activation across all cortical depths (strong main effect), whereas the effect of attention to the monocular stimulus only reaches (marginal) significance at the most superficial cortical depth. Still, the interaction between cortical depth and the attentional effect is highly significant at p < 0.001. ***C) The BOLD response was deconvolved across cortical depth using the model proposed by Marquardt et al.*** *(2018)*. This analysis showed a much flatter laminar profile of inferred activity. **D) *A plot of the laminar deconvolved responses.*** This analysis confirms the hypothesis that thalamo-cortical input processes and cortico-cortical feedback processes lead to different activity profiles across cortical depths. *\*\*\* p < 0.001, * p < 0.05, † p < 0.1*

The stimulation of the preferred eye of a voxel showed a large main effect across all cortical depths (F(1, 5) = 45.8, p = 0.0011 for left V1; F(1, 6) = 46.7, p = 0.005 for right V1), as well as an interaction with cortical depth (F(4,20) = 10.9; p = 0.0001 in left V1, F(4, 24) = 13.6; p < 0.001 in right V1): the effect of stimulation was larger in voxels closer to the CSF.

In contrast, the main effect of attention was only marginally significant for left V1 (F(1, 5) = 14.2, p = 0.013) and even non-significant for right V1 (F(1, 6) = 1.5, p = 0.26). However, we did find a highly significant interaction effect between attention and cortical depth in both left (F(4, 20) = 7.29; p = 0.0009) and right V1(F(4, 24) = 15.4; p < 0.001 in right V1). This indicates that although the overall effect of attention on the BOLD response in V1 is relatively weak, it is much stronger for voxels near the CSF (see Figure 5B).

Interestingly, there was no significant interaction between stimulation condition and attention (F(1, 5) = 0.08, p =0.78 for left V1; F(1, 6) = 0.23, p=0.65 for right V1), nor a three-way interaction between stimulation, attention and cortical depth (F(4, 20) = 0.08; p = 0.99 for left V1; F(4, 24) = 0.26, p=0.90 for right V1). In other words: the effect of attention on BOLD activity in ODCs was independent of whether the preferred eye of an ODC was stimulated or not. This is consistent with the notion that participants are not shifting their attention towards a specific eye.

To control for the effect of draining veins, where the BOLD signal in more superficial layers is contaminated by the signal of deeper layers, we used the linear laminar deconvolution approach developed by Marquardt and colleagues (Markuerkiaga et al., 2016; Marquardt et al., 2018). This deconvolution analysis also confirmed that the relevance of the monocular and binocularly presented stimulus has an effect *exclusively at superficial cortical depth* (at p < 0.05), whereas stimulation of the preferred versus non-preferred eye has *across all cortical depths*, with the largest effect in the *middle* layers (Figure 5C-D). This confirms a dissociation of neural activity related to thalamocortical input versus cortico-cortical feedback connections across cortical depth.

Also for the deconvolved laminar signals, there was no interaction between attention and stimulation condition (F(1, 5) = 0.407, p = 0.85 for left V1; F(1, 6) = 0.069, p =0.80 for right V1), nor a three-way interaction (F(4, 20) = 0.56, p =0.69 for left V1; F(4, 24) = 1.28, p = 0.30 for right V1). Again, this suggests that in our experiment, subjects are not able to shift their attention in an eye-specific manner.

### Decoding eye stimulation from moment-to-moment

Univariate results such as the ones reported above may overlook patterns in the data that multivariate decoding analyses do capture (Haxby et al., 2001; Haynes and Rees, 2006; Kamitani and Tong, 2005; Mur et al., 2009; Norman et al., 2006; O’Toole et al., 2007; Tong and Pratte, 2012). We therefore conducted an additional layer-specific decoding analysis to investigate the impact of attention on ODC patterns.

We constructed a Bayesian encoding model for monocular population coding, based on the model proposed by van Bergen and colleagues (van Bergen et al., 2015; van Bergen and Jehee, 2019). It assumes two independent neural populations coding for input of the left and right eye and voxels’ activity is then modeled as a weighted sum of these two populations and their (co)variance as a multivariate normal distribution, consisting of a weighted sum of independent voxel noise, shared voxel noise, and binocular population noise. The model was fit and evaluated per sampled time point, sampled at different cortical depths and cross-validated across runs (see methods section for details)

Stimulus eye-of-origin decoding was reliably above chance in all subjects (see Figure 6A for traces of the log_10_ Bayes Factor for all the runs of an individual subject), with volume-to-volume accuracies ranging from 66% – 84% for left V1 and from 68% – 91% for right V1.

**Figure 6.**
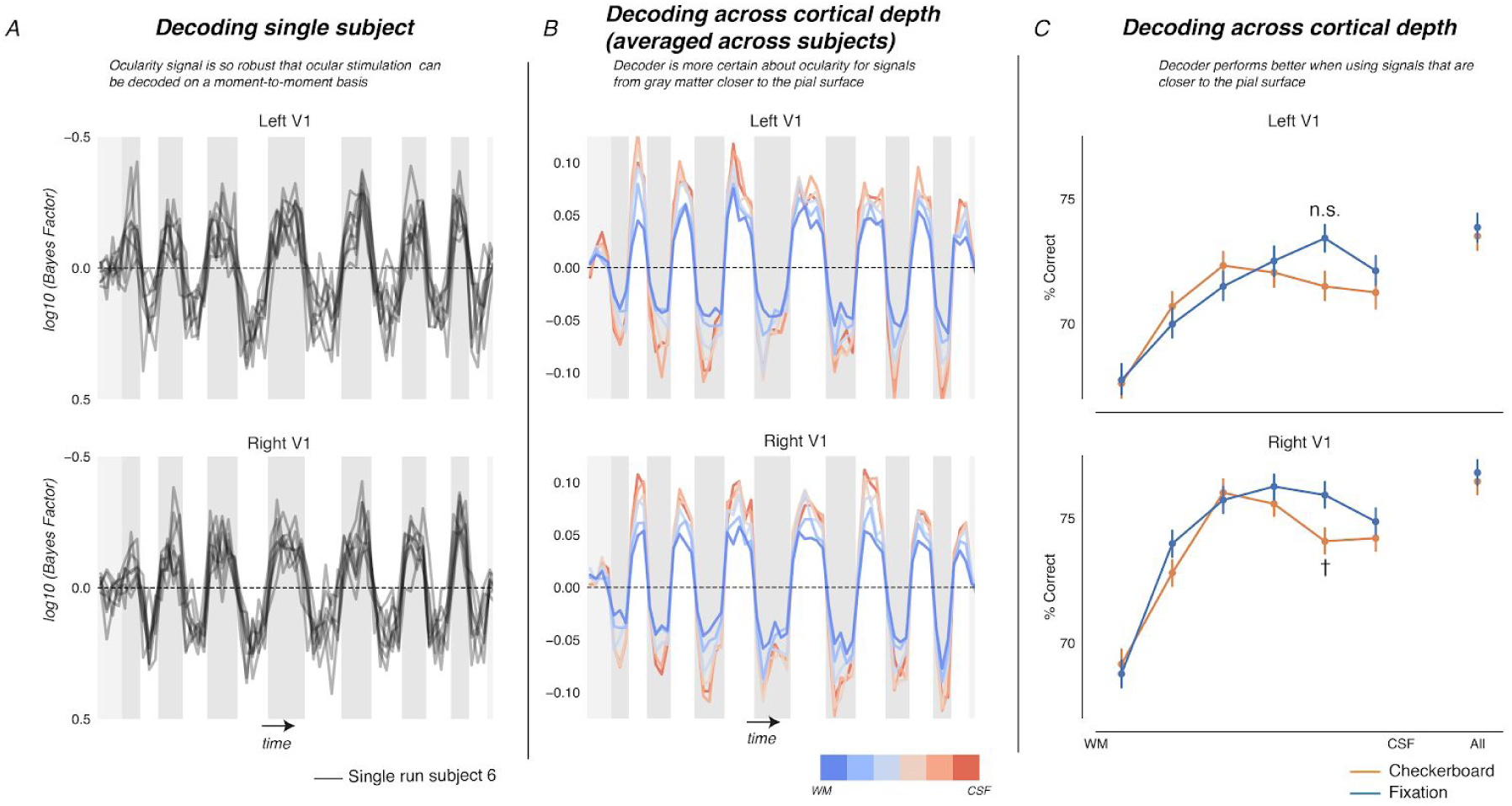
An inverted encoding model of neural ocular dominance populations can reliably decode the ocular stimulation condition on a moment-to-moment basis. ***A) Traces of the log_10_ Bayes Factor odds between left and right eye stimulation for all runs of a single subjects.*** Note how the original stimulus design can be read out from decoded, previously unseen data from V1 using the activation patterns of single fMRI volumes. ***B) Mean Bayes Factor trace across subjects, splitted across different cortical depths.*** Note how the model is more certain about its predictions for BOLD data that was sampled closer to the pial surface. **C) *Accuracy of the decoder for different cortical depths and attentional conditions.*** Error bars indicate bootstrapped standard errors of the mean (SEM). The decoder performs best at middle cortical depths. There are no significant differences between the two task conditions at any cortical depth. There is also no significant main effect or interaction effect related to the task conditions. *n.s. = not significant, † = p < 0.1*

Bayesian hierarchical model comparison showed, both for left and right V1, that the relationship between cortical depth and decoding accuracy is better explained by a quadratic model than a linear model (wAIC of −321.4 vs −257.54 for left V1 and −291.12 vs −217.78 for right V1). Furthermore, modelling suggests that highest decoding accuracy is achieved by using BOLD activity at a cortical depth of approximately 27 % cortical depth for left V1 (albeit with large uncertainty, 95% HPD: [-193%, 315%]) and approximately 45% for right V1 (95% HPD: [24%, 107%]). This cortical depth roughly corresponds to the layers at which our ODC patterns are strongest (Figure 4D). It is these layers where thalamocortical monocular fibers enter V1 and ocular neural activity patterns as measured by post-mortem techniques are most prominent (Adams et al., 2007; Adams and Horton, 2009; Dougherty et al., 2019).

There was no effect of attention condition on the accuracy of the decoder (F(1, 5) = 0.063, p = 0.81 for left V1 and F(1, 6) = 0.86, p = 0.38 for right V1), nor an interaction between the attention condition and cortical depth (F(5, 25) = 1.97, p=0.12 for left V1; F(5, 30) = 1.33, p = 0.28 for right V1). An apparent trend of *increased* decoder accuracy for the before-last cortical depth for *binocular* attention was not statistically significant (Figure 6C; t(6) = 2.10, p = 0.07 when both hemispheres are combined; also not significant for individual hemispheres). In sum, the pattern of results from the decoding analysis was consistent with the univariate results presented above: thalamocortical input is most strongly represented at deep and middle cortical depth, and spatial attention does not selectively gate information from a specific eye.

## Discussion

We applied 7 Tesla functional MRI to map ocular dominance columns (ODCs) in healthy human subjects, while participants focused their attention on either a monocularly presented stimulus or a binocularly presented fixation mark. Crucially, whereas earlier studies that mapped ODCs employed highly anisotropic “pencil voxels”, with slices 5.5 to 7.3 times thicker than the in-plane resolution (Cheng et al., 2001; Dechent and Frahm, 2000; Goodyear and Menon, 2001; Menon et al., 1997; Yacoub et al., 2007), for the first time, we mapped ODCs using isotropic voxels. This allowed us to study monocular BOLD activity in V1 along three dimensions: not only the 2D columnar location on the cortical surface, but also cortical depth (Kuehn and Sereno, 2018). The relationship between cortical depth and neural activity as measured by BOLD fMRI has the potential to be highly informative of brain function, since feedforward- and feedback-processes are believed to reside at different cortical depths (De Martino et al., 2018; Lawrence et al., 2019; Stephan et al., 2019). The combination of columnar and laminar sensitivity allowed us to measure the effect of thalamocortical input without manipulating stimulus properties, which are possibly confounded with other higher-order perceptual processes. The laminar patterns in our data align remarkably well with post-mortem tracing studies and electrophysiological recordings in rodents and non-human primates.

First, the effect of stimulating the preferred versus non-preferred eye, after correcting for draining veins, was most robust at deep and medium cortical depth, consistent with thalamocortical input into infragranular layer 6 and granular layer 4. Concordantly, the depth at which the decoding was most accurate was consistent with the granular layer and model comparisons show that its performance across depth is best described by an inverted-U pattern. Considering the effect of draining veins, where BOLD signals leak outwards towards the pial surface (Heinzle et al., 2016; Polimeni et al., 2018), we infer that the decoder was mostly relying on BOLD activity originating from deep and middle cortical depths. Interestingly, it is well-known that thalamocortical projections into V1 are most numerous in the granular layer 4c and this layer has so-far received most attention as the thalamocortical input layer of primary sensory cortex in laminar fMRI studies (Koopmans et al., 2010; Lawrence et al., 2019; Stephan et al., 2019). However, less well-known is that recent work in rodents has shown that neural activity in infragranular layer 6 also represents direct input from thalamus, with transmission latencies that are indistinguishable from those of layer 4C (Constantinople and Bruno, 2013). Furthermore, additional work in rodents has shown that thalamocortical projections in layer 6 tightly follow the columnar structure of layer 4 in rodent barrel cortex (Crandall et al., 2017). Intriguingly, Tootell and colleagues (1988) used radioactively labelled 2-deoxyglucose (DG)-uptake in post-mortem studies, to visualize ODCs in macaques and show that the largest DG uptake, after layer 4C, was to be found in layer 6. Similarly, in a recent study on binocular modulation of monocular neurons in layer 4 of V1 in macaques, Dougherty and colleagues (2019) found the largest proportion of monocular neurons in layer 4 and “layer 5/6” (their supplemental figure S1, panel D).

The effects of spatial attention on the laminar activation profile also fit earlier computational, anatomical and electrophysiological work. In our data, shifts of attention to the monocularly-presented stimulus features exclusively modulated BOLD responses near the pial surface. This is consistent with neurocomputational models of canonical microcircuitry where higher-order areas send feedback signals to supragranular layers of upstream cortical regions (Bastos et al., 2012; De Martino et al., 2018; Dumoulin et al., 2018; Stephan et al., 2019). It is also consistent with tracing studies that suggest that the majority of neurons in V2 that project to V1, project to supragranular layer 1 (Anderson and Martin, 2009; Rockland and Virga, 1989). Furthermore, other cortical depth-resolved fMRI studies have shown that the BOLD signal at superficial cortical depth is modulated by feedback processes in primary visual (; e.g., Muckli et al., 2015, Lawrence et al., 2019) and auditory cortex (De Martino et al., 2015). Finally, recent electrophysiological studies on attention and visual working memory in non-human primates, Van Kerkoerle and colleagues (2017) and Denfield and colleagues (2018) showed that attention correlates with increased top-down inputs to both infragranular layers and especially supragranular layers. An interesting question is why electrophysiological studies found effects of attention in infragranular layers, whereas our study (e.g., Muckli et al., 2015, Lawrence et al., 2019; as well as others) only find attentional effects at a cortical depth consistent with supragranular layers. Possibly, this difference in results is due to a significantly smaller change of neural firing in infragranular layers as compared to supragranular layers: in fact, Denfield and colleagues, in their electrophysiological work, found a significant effect in infragranular layers in only one-out-of-two attentional conditions (Denfield et al., 2018). We speculate that activity in the infragranular layers is representing mostly intracolumnar processing: neural activation from supragranular to infragranular layers (Bastos et al., 2012; Stephan et al., 2019) and that laminar fMRI is less sensitive to these short-range projections than the longer corticocortical feedback and thalamocortical feedforward projections. However, we can not exclude that the lack of an attentional effect at deeper layers is (partly) due to an inherently smaller BOLD response.

We did not find an interaction between the monocular stimulation of the preferred eye and task relevance of the monocular stimulus. This implies that although attentional processes influence activity in the supragranular layers of V1, they do not impinge on the ocular dominance activity pattern as measured by BOLD fMRI. That is, a shift of spatial attention, when directed to a monocularly presented stimulus feature, does not gate the lateral geniculate-cortical inputs to V1 based on their ocularity (eye-of-origin). Earlier work in the context of binocular rivalry has, however, shown that attentional processes can modulate monocular input streams (Zhang et al., 2011, 2012). One possible explanation for this apparent discrepancy in results pertains to the experimental paradigm used: possibly, attentional gating of the two eyes only happens when these send highly conflicting information, such as in standard binocular rivalry (Leopold and Logothetis, 1996; Levelt, 1965; Tong et al., 2006) and continuous flash suppression paradigms (Tsuchiya and Koch, 2005). In the paradigm used here, one eye was presented with a flickering checkerboard, whereas the other eye, at the same retinotopic location, was presented with a black background, as if this part of the stimulus was occluded in one eye. Although this difference between the two input streams does indeed represent a conflict, this conflict between monocular signals is much less salient than during full-fledged rivalry paradigms, where stimuli are purposely made highly incongruent, such as equal-contrast, orthogonal gratings of different colors (Haynes et al., 2005; Haynes and Rees, 2005; Zhang et al., 2012, 2011).

A second potential explanation of the lack of interaction between the input- and feed-back manipulation in our data, is that attentional gating of rivaling stimuli might not be resolved at the level of V1, but already earlier in the visual processing stream, at the stage of the lateral geniculate nucleus (LGN). Indeed, higher-order attentional effects and perceptual states can influence neural processing already at the level of LGN (Haynes et al., 2005; O’Connor et al., 2002; Wunderlich et al., 2005). Future work should elucidate the precise interplay between attention and ocular processing by measuring ODCs in V1 across cortical depth using UHF-fMRI during full-fledged binocular rivalry and simultaneously imaging ocular dominance layers in LGN (Zhang et al., 2010).

A recent 7T fMRI study by Lawrence et al. (2019) also investigated the laminar BOLD activation patterns of thalamocortical *bottom-up* and corticocortical *feedback* processes in V1. Lawrence and colleagues did not use columnar inputs structures, but different stimulus contrast levels to experimentally manipulate thalamocortical input. The feedback manipulation was also different, as a feature-based (as opposed spatial) attention manipulation was used: subjects had to report on one of two orthogonally oriented stimulus features of the same stimulus at the same retinotopic location. To correct for the increase of BOLD activity towards the pial surface due to draining veins, Lawrence et al. normalized (z-scored) the estimates of raw BOLD activity for their main results. This approach is comparable to our decoding approach, where the performance of the decoder is largely determined by the ratio between task-related activity and the variance of the BOLD signal. Indeed, Lawrence and colleagues found the largest effect of their input manipulation at medium cortical depth, as we did here. Their input also showed an inverted-U shape, consistent with our decoding results. However, Lawrence and colleagues interpret this U-shaped pattern as being evidence for thalamocortical input exclusively into the middle granular layer of V1. We propose that, since there is a significant effect of stimulus contrast near the white/gray matter border as well (both for standardized and raw data, their figure 3, supplement 5), it can not be excluded that this inverted U-shape is the result of thalamocortical inputs to both infragranular and granular layers. This speculation highlights the importance of modelling of the draining vein problem and/or taking this into account when interpreting raw BOLD effects across cortical depth (Heinzle et al., 2016; Marquardt et al., 2018; Polimeni et al., 2018). For the feature-based attentional manipulation, Lawrence et al. found the numerically largest (normalized) effect of attention at more superficial cortical depth, as we did here. They found a highly significant main effect of attention across *all* cortical depths and the interaction between cortical depth and effect size was non-significant (whereas we found a significant interaction effect at p < 0.001), suggesting that in their experiment, attention modulates all cortical layers equally. One explanation for this apparent discrepancy in results is that the resolution of both our anatomical (0.26 vs 0.51 ml) and functional data (0.34 versus 0.51 ml) was about 1.5-2 times higher than the data presented in Lawrence et al., which could have reduced BOLD signal blurring across laminae due to the increased fidelity of cortical surface reconstructions and reduced partial volume effects. Another explanation pertains to the properties of the task: in the experimental paradigm used by Lawrence et al., the orientation that needed to be attended by the subjects was explicitly relevant to the task. This is different from our task, where the relevance of the ocularity of the input signals is only implicit and of a different nature than the orientation of a stimulus. The relevance of orientation in the task of Lawrence and colleagues could have recruited additional processing of incoming signals, such as cross-orientation suppression. This processing could have happened via intra-columnar connections from the supra- to the infragranular layers. Future work should investigate different forms of attention, like spatial and feature-based attention using UHF-fMRI in the same scanning session, to determine whether these subtle differences in the laminar location of attentional effects are indeed due to fundamentally different mechanisms.

In conducting this study, we took the utmost care to tackle the wide range of methodological challenges that ultra-high field fMRI has to surmount. We stress here that, in our experience, extensive manual checking and correction of initial masking, unwarping, registration and segmentation are essential to make valid inferences about the relationship between cortical depth and BOLD activation patterns. We therefore included a very detailed methods section and shared all our analysis scripts^1^ and raw data^2^ to open source repositories. We think that such sharing of data and code is of crucial importance to understand what is and what is not possible with UHF cortical depth-resolved fMRI and to determine the level of methodological rigor needed to further this young and exciting field (Poldrack et al., 2019).

## Methods

### Participants

Seven participants were recruited from the Vrije Universiteit Amsterdam. They were aged between 21 and 39 years old. All participants gave informed consent, and procedures were approved by the ethical review board of the Vrije Universiteit Amsterdam.

### Apparatus

#### MRI Acquisition

All data from Amsterdam were acquired using a Philips Achieva 7 T scanner (Best, Netherlands). For the anatomical data, we used a volume transmit coil for excitation and a 32-channel head coil for signal reception (Nova Medical, MA, USA). For the functional acquisitions, data were acquired using two custom-built high-density 16-channel surface coils (total 32 channels) for signal reception (Petridou et al., 2013) and the standard NOVA coil for transmission.

MRI data from Beijing were acquired on a Siemens 7T Magnetom Scanner (Siemens Healthcare, Erlangen, Germany) with a 32-channel receive 1-channel transmit head coil (Nova Medical, Cambridge, MA, USA), from Beijing MRI Center for Brain Research (BMCBR). Gradient coil has a maximum amplitude of 70mT/m, 200 us minimum gradient rise time, and 200T/m/s maximum slew rate.

#### Stimulus presentation

Fully separated dichoptic stimulus presentation was achieved by use of prism glasses (Schurger, 2009), with a septum and a back-projection screen mounted inside the transmit coil. The quasicircular back-projection screen was 29cm in diameter to fit precisely into the transmit coil. Depending on the individual participant’s positioning, the resulting distance to the screen was between 12 and 15 cms. This caused variations in the maximal stimulus eccentricity (see below) across subjects, for which we corrected in our pRF eccentricity and size estimates. For some participants, we used positive-diopter lenses to aid their ability to maintain focus at this short distance. To minimize reflections inside the transmit coil it was covered in black cloth, and the septum was painted using matte black paint.

A ProPixx (VPixx technologies, Saint-Bruno, Canada) projector running at 1920×1080@120Hz was used for stimulus presentation, implementing a linearized lookup table. As the scanner environment is hostile to exact luminance output measurements of this custom setup the maximal luminance of the display is unknown. Based on the manufacturer’s data however, we estimate that it was over 500 cd/m2. Due to the shape of our projection screen and the distance between screen and projector, the width and height of the effective screen area were approximately 1400 by 700 pixels.

### MRI protocols

#### Anatomical data

For every subject, we collected both MP2RAGE (Marques et al., 2010) and MP2RAGE-ME (Caan et al., 2019) data. The MP2RAGE-ME (FOV: 204.8×204.8×200.6 mm; matrix size: 320×320×313; resolution 0.641 × 0.641 × 0.64 mm; TR_MP2RAGE_ = 6.246 s; TI_1_ = 0.67 s; TI_2_ = 2.7 s; FA_1_ = 7; TR_excitation,1_ = 0.0062 s; TE_1_: = 0.003 s; FA_2_ = 6°; TR_excitation,2_ = 0.0314 s; TE_2,1_ =; TE_2,2_ = 0.0155 s; TE_2,3_ = 0.02 s; TE_2,4_ = 0.0285) was collected because it offered both high-quality T1-weighted images, as well as images with T2* and PD-contrast (the four echoes of the second inversion images). These T2*-weighted images were useful for masking out the sagittal sinus from the gray-matter (GM)/white-matter (WM) segmentation (the sagittal sinus has the same image intensity as GM on standard T1w-images, which can cause severe problems for automated segmentation algorithms).

In addition, a standard MP2RAGE (FOV: 220 × 220 × 200.32 mm; matrix size: 352 × 352 x313; resolution: 0.625 × 0.625 × 0.64 mm;TR_MP2RAGE_ = 5.5 s; TI_1_ = 0.8 s; TI_2_ = 2.7 s; FA_1_ = 7°; FA_2_ = 5°; TR_excitation_ = 0.0062; TE = 0.003 s) was collected, because of its higher signal-to-noise and gray-matter/white-matter contrast and would later be combined with the MP2RAGE-ME images to aid the gray-matter/white-matter-segmentaiton(see *preprocessing anatomical data*).

To correct for residual B1+ effects in the MP2RAGE(ME), three DREAM B1-images (FOV: 224 × 224 × 224 mm; matrix size: 64 × 64 × 64; flip angles of 15, 30 and 45 degrees to cover a range of B1+ values were also collected.

Anatomical images from Beijing were collected with a T1-MP2RAGE sequence (0.7mm isotropic voxels, acquisition matrix = 320 × 320, GRAPPA acceleration factor = 3, TE = 3.05ms, TR = 4000ms, TI1 = 750 ms, Flip angle = 4 deg, TI2 = 2500 ms, Flip angle = 5 deg, FOV = 224 × 224 mm). Subjects used bite bars to restrict head motion.

#### Functional data

In Amsterdam, for the mapping of the ocular dominance columns, functional data was collected using the ultra-high resolution fMRI protocol described in Petridou et al. (Petridou et al., 2013). Specifically, we collected a 3D GRE-EPI protocol at a resolution of 0.7 mm isotropic with a volume acquisition time of 4 seconds (FOV = 120 × 120 × 22.1 mm; matrix size = 192 × 192; 30 slices; TR = 0.054 s TE = 0.027 s; FA = 20°; FOV, 120 × 120 × 21 mm (30 slices); echo planar factor, 27).

For the receptive field mapping paradigm, a higher temporal resolution was warranted, to reliably estimate PRF parameters from the design matrix (Dumoulin and Wandell, 2008). Therefore, for these runs, the number of shots and thereby spatial resolution was reduced to 1 mm (FOV = 160 × 160 × 34 mm; matrix size = 160 × 160; 34 slices; TR = 0.058 s; TE = 0.028 s; FA = 20°; echo planar factor 21). This protocol had a volume acquisition time of 2.7 seconds.

For both the 0.7 mm and 1.0 mm functional MRI protocols, we also collected an identical protocol, except with reversed phase-encoding blips (phase-encoding in the opposite direction). Those images would be used to correct for distortions due to B0 field inhomogeneities using the “TOPUP”-algorithm (Andersson et al., 2003). For the ocularity mapping paradigm, we collected 5 volumes with opposite phase-encoding. For the PRF mapping paradigm, we collected 8 volumes with opposite phase-encoding.

Submillimeter resolution (0.8 mm isotropic) fMRI data from Beijing were collected with a 2D GE-EPI sequence (31 slices, slice thickness = 0.8 mm, TR = 2000 ms, TE = 23 ms, FOV = 128 × 128 mm, image matrix = 160 × 160, GRAPPA acceleration factor = 3, FA = 80°, Acquisition Bandwidth = 1157Hz, phase encoding direction from Foot to Head). Immediately before each functional scan, five EPI volumes with reversed phase encoding direction (H to F) were acquired for distortion correction. Four functional scans were collected for the ocular dominance mapping experiment.

### Experimental Paradigm

#### Framing stimulus array

Both the PRF mapping paradigm, as well as the ocularity mapping paradigm were presented within a larger stimulus array (the *framing stimulus array*) that was designed to help the subjects comfortably fuse the input images of the left and right eye and thereby to prevent them from making unwanted eye movements. Before an experimental run started, the subject was able to move and rotate the left and right stimulus array with the response boxes, so that binocular fusion of the two stimulus arrays was as comfortable as possible. Specifically, the subject could translate, scale, and rotate the left and right stimulus array independently. The subject was instructed to make the framing stimulus array as large as possible, without any of its parts falling out of the visible area.

See figure 2B and 2D for a graphical representation of the framing stimulus array. It was built around a circular main stimulus aperture, with a diameter of 23-28 degrees-of-visual-angle. Its exact size depended on the shape of the subject’s head and the way the subject set up the stimulus at the beginning of the experimental run. On the outside of the stimulus aperture, an outer rim was presented with a width of 10% of the inner aperture (2.3 – 2.8 degrees-of-visual angle). This rim also contained four checkerboard squares (3×3 cells) with a size of 30% of the stimulus aperture, to annotate the cardinal orientations of the stimulus array, aiding in the binocular fusion of the subject.

The inner 5% radius of the stimulus array (approximately 1.15 – 1.4 degrees-of-visual-angle) was occupied by a fixation dot. The inner 33% of this dot was colored red/green, the middle 33% was occupied by a black circle, and the outer part of the fixation dot was a white ring. The colored dot in the middle of the fixation circle dynamically changed color according to an exponential distribution with a scale (λ) of 0.33, corresponding to a mean duration of 3 seconds. However, the distribution was left-censored, such that only durations of two seconds or longer could occur, which increased the effective mean duration of the fixation dot stimuli to 5 seconds.

#### PRF mapping paradigm

We used a standard “passing bar” PRF mapping paradigm (Dumoulin and Wandell, 2008) that was presented binocularly using the binocular stimulation setup described before (see Figure 2B for illustration). The passing bar had a height of 12.5 % of the main aperture size (approximately 2.875 – 3.5 degrees-of-visual-angle) and consisted of 2000 purple/cyan (50%) and green/blue (50%) overlaid gabor patches with a spatial frequency that was sampled from a uniform distribution between 0 and 30 cycles/degree. The Gabors patches were uniformly spaced across the passing bar and were dynamically and randomly repositioned according to a uniform distribution between 0 and 0.05 seconds. The size of the gabors was 0.75 degrees-of-visual angle. The Gabors were shifting phase at a frequency of either 3 (50%) or 12 Hz (50%). Finally, all the stimuli inside the passing bar were flickering together according to a square wave at a frequency of 12 hertz. The complete stimulus array was visually extremely salient and was designed to maximally modulate V1 activity and to optimize the efficiency of the PRF mapping paradigm. The power of the paradigm is reflected in the quality of the resulting retinotopic maps at a volumetric resolution of 1 cubic millimeter, without any smoothing and collected in just over 15 minutes.

Most subjects performed 3 runs of the PRF mapping paradigm (one subject performed 5 runs). Subjects were instructed to report changes of color (red/green) of the central fixation dot and ignore the passing bar stimuli.

The paradigm started with only a fixation dot for 24 seconds, then a bar pass from top to bottom in 30 seconds, a bar pass from top left to bottom right for 30 seconds, then a 12 second rest period with only the fixation dot and aperture stimulus array, then a 30 second bar pass from bottom to top, then a 30 second bar pass from bottom right to top left, then another 12 second rest period, then a 30 second right-to-left bar pass and 30 second bottom left to top right bar pass, another 12 second rest period and finally a 30 left-to-right and bottom left-to-top right bar pass. The total duration of the paradigm was thus 324 seconds (5 minutes and 24 seconds). At the end of an experimental run, 8 different and angularly equidistant bar passes were shown to the subject. The paradigm ended with a 24 second period with only the fixation dot and stimulus array.

#### Ocular Dominance Mapping Paradigm

For the ocular dominance mapping paradigm, we used the same general stimulus array as the PRF mapping experiment, but the moving bar was replaced with a circular rotating and flickering checkerboard (see Figure 2D for illustration). The flickering checkerboard fitted exactly in the inner aperture (23 – 28 degrees-of-visual-angle), but was masked by a raised cosine, reducing the contrast for the outer 20% radius of the checkerboard.

The cells within the checkerboard had a width and height of 2 degrees-of-visual-angle. The contrast of the checkerboard was inverted (black cells become white and vice versa) according to a square wave with a frequency of 8 Hz. The Checkerboard was also rotating at a frequency of 0.66 rotations per second. The direction of the rotation (clockwise versus counter-clockwise) was changed once every 5 seconds on average, according to the exact same left-censored exponential distribution as the color of the fixation dot.

Importantly, unlike the framing stimulus array, the checkerboard was only presented in *one eye at a time* (monocular stimulation). It was first presented to the left eye for 12 seconds, then the right eye for 12 seconds. Then the paradigm continued with subsequent monocular stimulation for 16, 20, 24, 20, 16 and again 12 seconds. This particular block design with immediate changes in ocularity was chosen to wash out any non-eye-specific effects by oversaturating their resulting BOLD-response (Moon et al., 2007). Before and after the monocular stimulation, an empty stimulus array was presented for 12 seconds. Thus, a single experimental run had a duration of 264 seconds (4 minutes and 24 seconds). Two subjects performed 10 runs of the binocular mapping paradigm, the other 5 subjects performed 8 runs.

On uneven experimental runs, subjects were verbally instructed to report *changes in color of the fixation dot* with a response button box, thereby focusing their attention on the *binocularly* presented part of the stimulus array. On even experimental runs, the subjects were verbally instructed to report *changes in the direction of the rotation of the checkerboard*, with their response button box, thereby focusing their attention on the part of the stimulus array that was *monocularly* presented.

### Data Analysis

#### Structural preprocessing

To study the 2D (cortical location) and 3D properties (laminae) of the functional organization of V1, precise delineation of the inner and outer border of the cortical surface is of the utmost importance (Polimeni et al., 2018; Waehnert et al., 2014). Therefore, we spent a lot of resources in a hybrid approach of automatic and manual segmentation of gray and white matter and CSF, as well as the subsequent reconstruction of the cortical surface. Because the correspondence between locations on the inner and outer surface of the cortical sheet was central to some of our research questions, we mainly opted for the surface inflation-approach of Freesurfer (Polimeni et al., 2018) in most of our analyses, rather than level set-based approach such as implemented in CBS tools (Bazin et al., 2014; Huntenburg et al., 2018). However, for some analyses we did use the latter approach to estimate cortical depth more precisely on a voxelwise basis. Furthermore, we used a combination of multiple software packages to preprocess and segment the anatomical data. The two end goals of the structural processing workflow were (a) a cortical reconstruction by Freesurfer represented as a white matter and pial mesh, consisting of points and triangles (Dale et al., 1999), (b) a white matter and pial surface reconstruction by CBS-tools, represented by two level sets (Bazin et al., 2014; Waehnert et al., 2014). To get the best possible segmentations with minimal manual intervention, we found it very helpful to combine the results of multiple segmentation algorithms as a “wisdom-of-the-crowd”-approach. See below for more details. We used the in-house developed python package *pymp2rage* (de Hollander, 2018) to estimate the T1-UNI image (Marques et al., 2010) and T1 maps of the MP2RAGE and MP2RAGE-ME data. We used the B1+ maps to correct for any residual B1+ bias fields (Marques and Gruetter, 2013). Furthermore, we also estimated T2*/R2*-maps from the four echoes of the MP2RAGE-ME data using ordinary least-squares in log signal space.

The next part of the structural preprocessing pipeline is implemented in a Docker image that can be found at https://github.com/VU-Cog-Sci/mp2rage_preprocessing. The T1-UNI image of the M2RAGE-ME was registered with 6 degrees-of-freedom (rigid body transform) to the T1UNI of the M2RAGE protocol using FLIRT (Jenkinson et al., 2002) and then resampled using windowed Lancos sinc-interpolation as implemented in ANTS (Avants et al., 2014). We corrected the mean second inversion (INV2) for bias fields using N4BiasFieldCorrection (Tustison et al., 2010), distributed with ANTs 2.2.0 and used this as input to FSL’s brain extraction tool (Jenkinson et al., 2012; Jenkinson, M., Pechaud, M., & Smith, S., 2005). The skull-stripped average INV2 was then used as input to AFNI’s AutoMask mask with a *clfrac*-parameter setting of 0.5. to estimate the background noise level and remove voxels from the mask that only contain noise.

Then, we took the averaged image of the two T1UNI images, the two INV1 and INV2 images, as well as the T1 maps and used them as input to the *MP2RAGE skull strip* and *MP2RAGE dura estimation* modules of CBS tools (Bazin et al., 2014) as wrapped by Nighres (Huntenburg et al., 2018) to make masks of the dura mater as well as the skull. The probabilistic map of the dura as estimated by Nighres was thresholded at 0.8 to obtain a discrete mask of the dura. The dura mask was then dilated by 2 voxels, excluding voxels that were manually or automatically labeled as brain by the BET algorithm.

Lastly, we made a very crude manual mask of the sagittal sinus using large voxel drawn using FSLEyes (McCarthy, 2019). All voxels within this mask were then thresholded based on their estimated R2* (1/T2*) value (voxels inside the sagittal sinus have very high R2* values due to susceptibility effects). The precise threshold was manually determined but was on average approximately 1/20. This semi-automatic approach yielded a very precise mask of the sagittal sinus.

The dura and skull masks generated by Nighres, as well as the noise voxel mask estimated by AFNI as well as the mask of the sagittal sinus were set to zero in the averaged T1UNI image. The masked T1UNI image was then used as input to the structural preprocessing module of fmriprep (Esteban et al., 2019).

In that pipeline, the T1-weighted (T1w) image was corrected for intensity non-uniformity (INU) with N4BiasFieldCorrection (Tustison et al., 2010), distributed with ANTs 2.2.0, and used as T1w-reference throughout the workflow. The T1w-reference was then skull-stripped using antsBrainExtraction.sh (Avants et al., 2008), using OASIS30ANTs as target template. Brain surfaces were reconstructed using freesurfer’s recon-all (Dale et al., 1999), and the brain mask estimated previously was refined with a custom variation of the method to reconcile ANTs-derived and FreeSurfer-derived segmentations of the cortical gray-matter of Mindboggle (Klein et al., 2017). Spatial normalization to the ICBM 152 Nonlinear Asymmetrical template version 2009c (Fonov et al., 2009) was performed through nonlinear registration with antsRegistration (ANTs 2.2.0), using brain-extracted versions of both T1w volume and template. Brain tissue segmentation of cerebrospinal fluid (CSF), white-matter (WM) and gray-matter (GM) was performed on the brain-extracted T1w using FAST (Zhang et al., 2001). The fmriprep preprocessing workflow yielded a cortical surface representation in Freesurfer format in “*hires*”-mode, with an average edge length of 0.55 mm in the white matter surface and 0.66 at the pial surface and that was used in further analyses.

The masked T1UNI, T1map and skull and dura masks of Nighres were also used as input to the MGDM segmentation algorithm (Bazin et al., 2014; Bogovic et al., 2013), as wrapped by Nighres (Huntenburg et al., 2018). Then, to create a level set-based cortical surface representation, the *voxelwise* GM/WM/CSF-segmentations of FAST (Zhang et al., 2001), Freesurfer (Dale et al., 1999), MGDM (Bazin et al., 2014; Bogovic et al., 2013), as well as manual “correction masks” (see below) were averaged and used as input to the CRUISE cortical surface reconstruction algorithm (Han et al., 2004), as implemented in Nighres (Huntenburg et al., 2018). The manual masks were weighted 5 times more than the automatic segmentation algorithms. This “wisdom of the neuroimaging software crowd”-approach led, in our experience, to much more precise cortical surface reconstructions than using only the MGDM (or any other) segmentation as input to the CRUISE cortical surface reconstruction algorithm.

After these two pipelines were run, both the pial and white matter surface meshes of Freesurfer and the levelset representations of CBS-tools were inspected in FSLEyes (McCarthy, 2019). Any missegmentations were annotated with manual “correction masks”, after which the entire preprocessing pipeline was run again, using these manual masks as additional input. We usually needed 3-4 of these iterations to get satisfying cortical surface reconstructions.

When the Freesurfer cortical surface reconstruction was sufficient, we made 6 additional surfaces between the pial and white matter surface using the “equivolumetric layering” formulas described by Waehnert et al (Waehnert et al., 2014) as implemented by Konrad Wagstyl in his “Surface Tools”-package^3^.

#### Functional preprocessing

Functional data was minimally processed and extra care was taken to not smooth the data by resampling the data only once. The preprocessing workflow was implemented in nipype (Gorgolewski et al., 2011) and can be found on GitHub^4^. It was identical for the PRF and ocular mapping paradigm.

The following steps were performed. First, each run was independently motion-corrected using linear rigid-body registration and a normalised correlation cost function towards the mean of the run with the MCFLIRT algorithm as implemented in FSL (Jenkinson, 2002). The same motion-correction was applied to the reverse phase-encoded data. Then, average images for both time series were constructed, where we used only the middle volumes of the task data, such that the unwarping algorithm would use the same number of volumes for the main task and reverse phase-encoded data. These two averages images were bias field corrected using N4BiasFieldCorrection (Tustison et al., 2010) after which they were registered towards each other (unwarped) following the “TOPUP” unwarping algorithm described by Andersson et al. (Andersson et al., 2003), as implemented in the *qwarp* program of AFNI (Cox, 1996) using median filtering across 1 mm and a minimum patch size of 5 mm. Then, the unwarp field was applied to the averaged task data (this time using all volumes in the run), using Windowed Lancos Sinc interpolation.

The averaged task data was then registered to the T1-weighted (T1UNI) anatomical data using the Gray/White-matter segmentation of MGDM as input to the boundary-based registration (BBR) algorithm as implemented by FSL’s FLIRT (Greve and Fischl, 2009; Jenkinson, 2002), with 9 degrees-of-freedom. We chose the 9 degrees of freedom, with additional scaling across the axes in addition to rigid body transformations, to allow for some flexibility in correcting any global residual B0 distortions. Then, the *MeasureImageSimilarity* function of ANTS with a Mutual Information cost-function was used to detect which of all the task data runs that was most similar (registered best) to the T1-weighted data. This was then used as a ‘reference BOLD image’ to which all the other runs were registered once more.

This last step turned out to be helpful, since for most subjects 1 or 2 runs were completely misregistered (due to the very small field-of-view that hinders many automatic registration algorithms) and it increased the overlap between subsequent task runs was often not sufficient.

Finally, for every volume of every run, (1) the motion-correction affine matrix, (2) the distortion unwarp field, (3) the registration to the T1-weighted anatomical image and (4) the refined registration to the best-registered reference BOLD image were all concatenated, so that the resampling of every individual volume in the timeseries to anatomical space could be done in one single Windowed Lanzcos Sinc interpolation step, as implemented in ANTS (2.2.0). This yielded an unsmoothed volume time series at a resolution of 0.7 mm isotropic, very well-aligned with the T1-weighted anatomical data.

We then used the white-matter/gray-matter segmentation of MGDM to create aCompCorr time series that can be used to regress out spatially correlated structured noise. ACompcorr components are time series based on projections of raw data on the largest components of a PCA decomposition of the signal in an eroded mask of the CSF and white matter, that is believed to represent structured (physiological) noise (Behzadi et al., 2007). We also extracted the translations and rotations of the MCFLIRT motion correction (Friston et al., 1996), as well as Framewise Displacement (Power et al., 2012) and DVAR (Power et al., 2014; Smyser et al., 2010). The functional data were also high pass-filtered by subtracting the output of a Savitzky-Golay filter with a polynomial order of 3 and a window length of 120 seconds (Savitzky and Golay, 1964). The resulting time series were then divided by the average image intensity of the corresponding voxels and divided by 100, to get a “percent signal change” time series.

For the purpose of fitting the PRF model, we then sampled the raw BOLD time series to surfaces vertices by taking the mean across all cortical depths for the PRF model using the *SampleToSurface* function of Freesurfer (Dale et al., 1999) and using trilinear interpolation.

#### PRF fitting

For fitting the PRF model, we sampled vertex-wise time series across the entire depth of the vortex. We did so because the retinotopic locations of neurons at the same 2D cortical location across cortical depth are very similar (Fracasso et al., 2016) and it allowed us to pool data across voxels, increasing the signal-to-noise ratio and the quality of the estimates of the retinotopic location of different 2D locations in V1. To further increase the signal-to-noise ratio of the data, we also linearly projected the DVAR time series, the Framewise Displacement, translation and rotation-parameters from the motion correction, as well as 6 anatomical CompCorr-components out of the raw signal (Power et al., 2014). Then, we estimated a standard Gaussian PRF model (Dumoulin and Wandell, 2008) with 3 parameters: angle and eccentricity (equivalent to x/y-coordinate of the Gaussian) and a size (standard deviation of the Gaussian). We used the popeye Python package (DeSimone et al., 2016) to construct hypothetical time series for a large range of parameters (25 eccentricities × 32 angles × 25 sizes). We then correlated these hypothetical time series with the actual time series for every vertex, sampled across its entire depth and took for every vertex the parameter settings with the highest correlation. This “grid search” approach already yielded a relatively coarse retinotopic map of V1, which were used as a starting point for a gradient descent optimization algorithm as implemented in PopEye (DeSimone et al., 2016). The resulting parameters were then plotted back onto the surface using Pycortex (Gao et al., 2015). We used an (arbitrary) R2 threshold of 15% within-set explained-variance explained to mask out very noisy vertices. Then, we manually delineated the left and right V1/V2-border on the polar angle maps of every individual subject. The borders of V1 and V2 were defined by visual angle phase reversals at the dorsal (lower vertical meridian of the visual field) and ventral (upper vertical meridian of the visual field) banks of the calcarine sulcus (Wandell and Winawer, 2011).

#### Estimation of ocularity maps

To estimate ocular dominance of cortical locations, we estimated a simple general linear model (GLM) using nistats (“Nistats,” 2019) on the preprocessed, voxelwise time series. The GLM consisted of one regressor representing stimulation of the left eye, convolved with a standard HRF, and a second regressor represented stimulation of the right eye, convolved with a canonical HRF. We also added 6 nuisance regressors, which were the 6 components that explained most variance of a principal component (PCA) decomposition of all the 14 collected nuisance regressors: volume-wise DVAR, framewise displacement, six translation and rotation parameters provided by the motion-correction algorithm, as well as six anatomical CompCorr-regressors. We opted for this PCA-approach because the nuisance regressors were highly correlated, and the total number of volumes (degrees-of-freedom) of the ODC data was low due to our 4-second TR.

We then fit parameter estimates for the average BOLD activation for left and right eye stimulation, as well as the contrast for left > right eye stimulation for every run separately, as well as over all runs combined. These parameter estimates were normalized for their variance, resulting in corresponding z-maps.

We then used the cortical depth estimates for every voxel provided by the CRUISE cortical surface level sets (Bazin et al., 2014; Han et al., 2004; Huntenburg et al., 2018) to assign voxels to 5 bins of equivolumetric cortical depth. We used those bins to estimate the effects of cortical depth on the height of the BOLD response in different task conditions, as well as its interaction with stimulation condition and/or attentional condition. We also applied the linear laminar deconvolution approach developed by Marquardt and colleagues (Markuerkiaga et al., 2016; Marquardt et al., 2018). Here, the activation of 5 different bins was modeled as a linear combination of 5 underlying neural populations. For example, the BOLD response at deepest layer is assumed to be a result of BOLD activation due to neural activation only in the deepest layers, whereas the 2nd-to deepest layer is assumed to be a linear sum of both BOLD responses to the neural activity in the deepest layer and the 2nd-to-deepest layer, etc.

The “left > right” contrast z-map was also sampled to 6 equivolumetric cortical surfaces as defined by Freesurfer. This was done for the analysis of the spatial frequency of the ocularity profiles on the cortical surface, as well as to gauge the consistency of the ocular dominance columns within subjects across runs. To not induce any spatial smoothing, *nearest neighbor* interpolation was used for this final interpolation step.

After estimating the z-maps, they were visually inspected on the surface, with the V1/V2-masks overlaid. All z-maps were visually consistent with ocular dominance column patterns: except for one hemisphere in one subject, they showed relatively large changes in z-value (2-4) over relatively short amounts of geodesic distance (∼1.5 mm). The left hemisphere of subject 1, however, showed a very large right-sensitive blob of activity in parafoveal V1. Since these patterns did not qualitatively align with ocular dominance patterns, this hemisphere was excluded from all further analyses. This is why in almost all analysis there was one more right hemisphere than left hemispheres. After visual inspection of the z-maps, we tested them for robustness by correlating the z-map of the first half of the session with the second half of the session. For one subject (subject 5), we found no consistent pattern. Both hemispheres of this subject were excluded from all further analyses.

#### Spatial frequency estimation on the surface

To estimate the spatial frequency of the ocularity maps, we applied 2D Gabor filters (Forsyth and Ponce, 2015) to the ocularity maps after resampling them to a flattened cortical surface. We used Freesurfer’s *tksurfer* to cut out V1 from the rest of cerebral cortex and used Freesurfer’s flattening algorithm (*mris_flatten*) to convert the 3D vertex coordinates to 2D plane coordinates, minimizing any distortions in their distance matrix. We then resampled the ocularity z-maps from their flattened 2D coordinates to a gridded plane with a resolution of 0.35 mm using linear interpolation. We then convolved this plane with Gabor patches with a frequency ranging from 1 mm/cycle up to 10 mm /cycle in 50 logarithmically spaced steps and 16 equally space orientations. Then, for every vertex, this yielded an estimate of its power for a specific frequency and orientation. We assigned a “main frequency” to every vertex by finding the frequency with the maximum amount of power, summed over all orientations. We then plotted the distribution of main frequencies across vertices and estimated its mode (most common frequency). We repeated this procedure across multiple cortical depths, so we could investigate any shift in main frequency and power across cortical depth.

#### Inverted encoding model

To estimate the amount of information that the entire pattern of activity in V1 contained, we used an adaptation of the inverted encoding model described by van Bergen et al. (2015). Specially we assumed that the BOLD acitvity in each vertex in V1 could be explained by a weighted sum of a “left eye” and “right eye” neuronal population. This weight matrix W (as many rows as vertices and 2 columns -one column represent the left eye, the other the right eye-) was estimated by solving the linear system

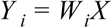

Where *Y* _*i*_ is the time course in vertex i and X is a design matrix with two columns, indicating which eye was stimulated at a given frame (either 0 or 1, shifted by 4 seconds in time to correct for hemodynamic delay). To regularize the weights matrix, we used ridge regression with a prefixed alpha-parameter of 1.0 (Hoerl and Kennard, 1970; Nunez-Elizalde et al., 2019), corresponding to a Gaussian prior with standard deviation of 1 on the weights. This regularizes the found weights towards more plausible values near 0.

Using the fitted weight matrix of the vertex-wise GLM, we fitted a regularized multivariate normal to the unexplained residuals *Y* _*i*_ − *W*_*i*_ *X* :

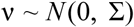

Where the covariance matrix Σ was defined as a weighted sum weighted of the diagonal covariance matrix *I* ∘ ττ ^*T*^ and a perfect correlation matrix ττ ^*T*^ (Ledoit and Wolf, 2003), as well as the estimated variance of the neural populations, σ^2^ (van Bergen et al., 2015), and finally the empirical covariance matrix Σ_*sample*_ (van Bergen and Jehee, 2019):

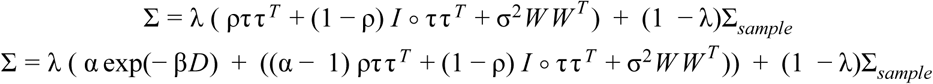

The weight matrix W was estimated with ridge regression (see above), λ represents the amount of regularization and was set to 0.9 based on cross-validation (performance gains compared to λ of 0.1, 0.5 or 1.0 depended on the amount of vertices used, but were in the order of 5% percentage point classification accuracy), Σ_*sample*_ is directly estimated from the data (cross product of data matrix) and ρ, τ, and σ^2^ are optimized using gradient descent. We implemented the model in Tensorflow (Abadi et al., 2016), which calculated the gradients of the parameters with respect to the parameters automatically (Bartholomew-Biggs et al., 2000; Corliss et al., 2002), thereby drastically improving the speed of the optimization process. To further speed up the optimization process, we selected a subset of 400 vertices in the V1 mask that showed the highest explained variance (R^2^) in the GLM to be used for the residual model.

After fitting the residual model, we estimated the maximum-a-posteriori (MAP) stimulus for unseen data D by inverting the likelihood model using an informed stick prior that only allowed either activation of the left eye with 1 *or* the activation of the right eye with 1:

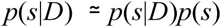

where

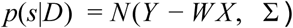

We then used the (log) Bayes Factor of the probability of a left versus right eye stimulation as a measure of decoded-eye and its uncertainty:

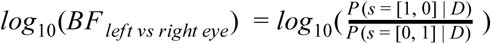

The log_10_(BF) represented the logarithm of the odds that the left or right eye was stimulated. When log_10_(BF) was above 0, the model predicted left eye stimulation, whereas if log_10_(BF) was below 0, the model predicted right eye stimulation.

We used a two-leave-out cross-validation scheme: For every fold, two runs were left out for validation purposes: one run with attention on the fixation cross and one run with attention on the checkerboard.

#### Linear versus quadratic model estimation

For two variables, namely total spectral power and decoding accuracy, we wanted to see if they increased or decreased linearly with cortical depth, or that they showed a clear peak at a specific cortical depth. To do this, we performed Bayesian model comparison. Specifically, we fitted two hierarchical linear model using pymc3 (Salvatier et al., 2016) and its implementation of the NUTS sampler (Hoffman and Gelman, 2014).

The dependent variable (decoding accuracy and spectral power) was first modeled as a linear function of cortical depth:

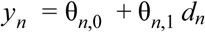

and then also as a quadratic function of cortical depth

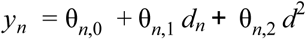

Where *y*_*n*_ were all observations for subject *n, d* is a corresponding vector of cortical depths and θ_*n*_ is the parameter vector for subject *n* that was estimated.

The subject-wise parameters were modeled as coming from a Gaussian group distribution mean μ and standard deviation σ :

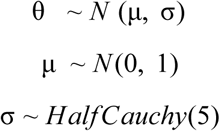

We used the state-of-the-art Watanabe–Akaike information criterion to do Bayesian model comparison between the linear and quadratic models (Gelman et al., 2014; Watanabe, 2013). Furthermore we, estimated the posterior of the peak of the quadratic function using the formula 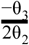 and calculated its 95% confidence interval using the highest posterior density approach (Kruschke, 2014).

We collected 4 chains of 2000 samples each, after 1000 tuning (“burn-in”) steps.

## Supplementary materials

### S1: The effect of cortical folding on effective resolution on the surface

In a recent paper (Kay et al., 2019), Kay and colleagues showed that the cortical surface reconstruction as implemented in Freesurfer, where every vertex in the white matter surface has a corresponding vertex at the pial surface, can lead to biased sampling at outer cortical depth. Due to dilations and compressions of the outer cortical surface compared to the inner surface at locations with high curvature, the number of vertices per surface area can increase/decrease compared to inner cortical depth.

We set out to test whether the effect described by Kay et al. reproduce in our high-resolution data and to which extent they could bias our spatial frequency results. We re-implemented the visualization technique they proposed (their Figure 4) to visualize distortions of our 0.7 mm isotropic voxels on the surface (Figure 7A). Indeed, we find some distortions, especially around areas of very high curvature. These distortions might bias the spatial frequency estimations.

**Figure S1.1:**
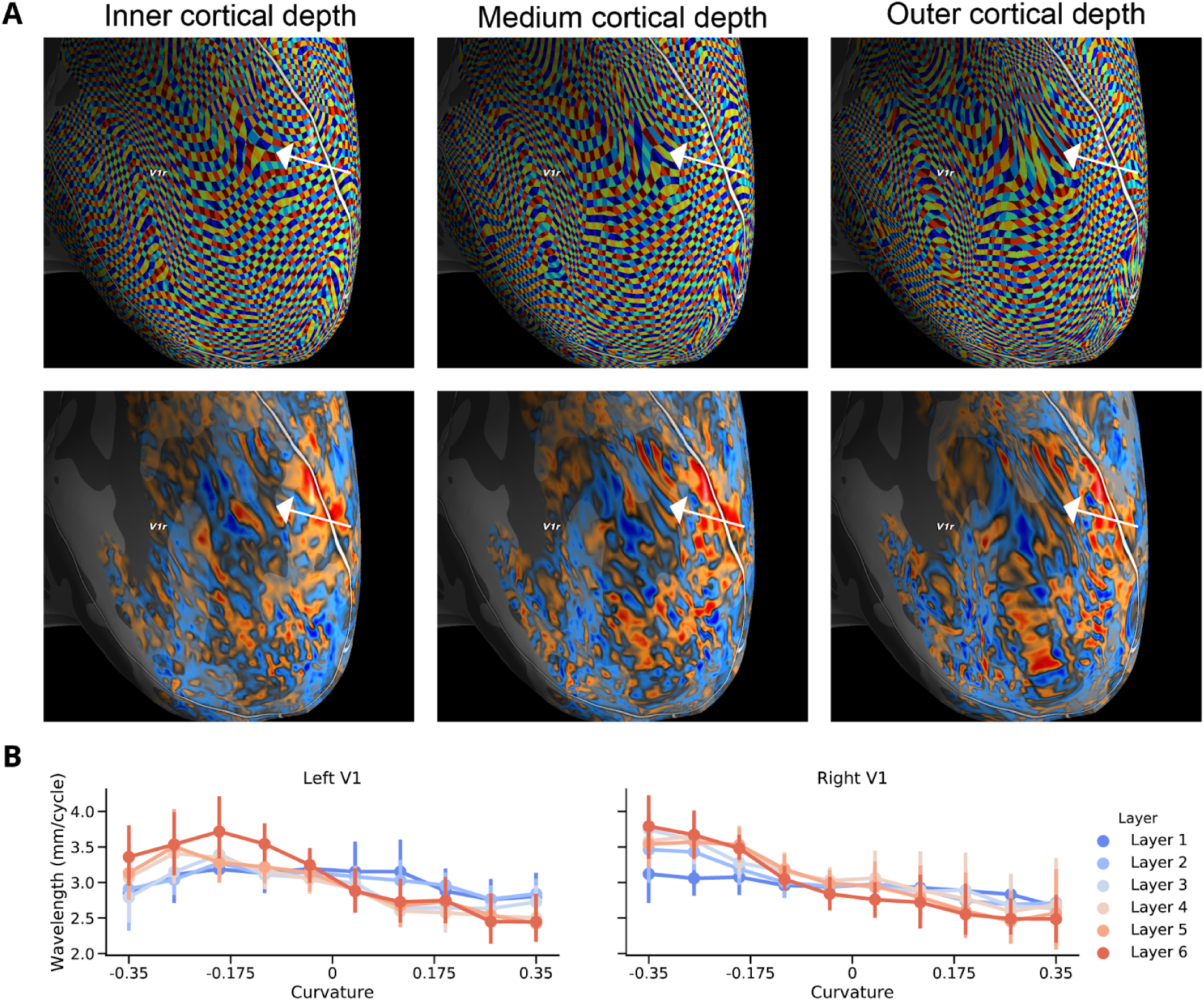
Strong curvature of the cortical sheet can lead to distorted spatial frequency estimates. **A)** Visualization of how original volumetric data at 0.7 mm isotropic resolution is sampled to the cortical surface across different cortical depths (Kay et al., 2019). Areas of high curvature lead to ‘distorted’ sampling and the white arrow annotates such an area. Note how the distortion gets larger at outer cortical depth. Also note how most of V1 does not seem affected by such distortions. **B)** A plot of the estimated wavelengths for vertices of different curvature. Especially for outer cortical depths (e.g., Layer 6), there is a bias towards longer wavelengths for negatively curved patches of cortex (sulci) and a bias towards shorter wavelengths for positively curved patches of cortex (gyri). However, when there is no significant curvature, wavelengths are the same across cortical depth. Furthermore, the average estimated wavelength across all curvatures are not statistically different from the unbiased estimates at completely flat patches of cortex.

Therefore, we plotted the modes of the spatial frequency distributions as a function of curvature (Figure 6B). Indeed, for outer cortical depths, we found larger spatial frequencies for negative curvature (sulci) and smaller spatial frequencies for positive curvature (gyri). When we performed a 2-factor ANOVA on the main spatial frequency with the factors depth (6 levels) and curvature (10 equally-spaced bins between −.35 and .35), we found a main effect of curvature (F(9, 45) = 13.77, p < 0.001 for left V1, F(9, 54) = 17.09, p < 0.001 for right V1), as well as an interaction effect between curvature and depth (F(45, 225) = 2.16, p=0.0001 for left V1 and F(45, 270) = 2.52, p<0.001 for right V1). However, there was no main effect of depth (F(5, 25) = 0.22, p=0.95 for V1 and F(5, 30) = 1.04, p = 0.41).

The main effect of this ANOVA is consistent with the undersampling of highly curved sulci. We suspect that the interaction effect of curvature are due to distortion of the vertex-to-vertex distances as the cortex gets flattened. This effect is worsened for outer cortical depths, because we used the same 2D coordinate system across all depths, which is based on the flattened *white matter* surface. The biases of these distortions can be ruled out by only analyzing areas of very low curvature. When we only analyze vertices with very low curvature between −0.05 and 0.05 (corresponding to a sphere with a radius of 20 cm or larger), no effect of depth on spatial frequency could be found in a one-way ANOVA (F(5,25) = 0.88, p =0.50 for left V1 and F(5, 30) = 0.45, p = 0.81 for right V1), just as in the analyses including all vertices in V1. Furthermore, the main spatial frequencies that are found in this analysis were not statistically different from those when we used vertices of all curvatures (F(5, 25) =0.26, p=0.93 for left V1; F(5, 30) = 0.93, p=0.47 for right V1). The finding that there is no difference between using only the most uncurved vertices and all vertices, suggests that the positive curvature and its dilation of the vertex in the gyri effectively ‘compensates for’ the negative curvature and compression in the sulci.

### S2: Relationship between spatial frequency of ocular map on the surface and retinotopy

By sampling Ocular Dominance Columns to the cortical surface, they can be compared to other features of visual processing that are organized topographically along the surface, such as the retinotopic representation of the visual field in V1 (Wandell et al., 2007; Dumoulin and Wandell, 2008; Wandell and Winawer, 2011). Therefore, in addition to our ocular mapping paradigm, we also performed a retinotopy mapping paradigm to delineate V1 and its retinotopic organization at an individual basis (Dumoulin and Wandell, 2008). This allowed us to relate properties of the ocular dominance columns to their position in the visual field. Several relationships between the properties of ODCs and their location in the visual field have been found in post-mortem studies in humans and other primates: (a) ODCs are slightly wider near the V1/V2-border (the upper and lower vertical meridian); (b) the columns from the contralateral eye become wider and thus more dominant at more eccentric parts of the visual field (specifically above 15 degrees) and (c) at about 15 degrees eccentricity, at the blind spot, there are no ocular dominance columns, and an oval area, corresponding to the size of the blind spot in visual space, is completely dominated by the ipsilateral eye (Adams et al., 2007; Adams and Horton, 2009). We tried to replicate all three findings in our in vivo fMRI data.

We used the estimated angle in the visual field for each vertex location as a measure of its distance to the V1/V2-border. Cortical locations at the V1/V2-border have a visual angle close to either −90 (upper vertical meridian, ventra V2) or +90 degrees (lower vertical meridian, dorsal V2). We divided the visual field in 6 equal angular bins ([0, 60], [60, 120], [120, 180], etc.) and tested whether there was a difference in wavelength across cortical locations corresponding to these six bins. Our hypothesis was that near the two vertical meridians, the spatial wavelengths of the ocularity pattern should be larger. However, one-way ANOVAs showed no significant effect of visual angle on wavelength for left V1 (F(5, 25) = 2.09, p=0.099), nor right V1 (F(5, 30) = 0.11, p = 0.99). Therefore, we could not replicate the pattern of wider ODCs near the V1/V2-broder described in post-mortem literature.

To test whether the contralateral eye becomes more dominant above 15 degrees eccentricity, we split vertices with a significant ocularity response (abs(z) > 1.96, p<0.05) in vertices with more and vertices with less than 15 degrees eccentricity. A one-way ANOVA showed no significant effect of eccentricity on wavelength of the ODC map in either left (F(1, 5) = 0.75, p = 0.42) or right V1 (F(1,6) = 0.29, p = 0.61). Thus, the pattern that is observed in human post-mortem data, where contralateral dominance is increased at high eccentricities did not replicate in our data.

### S3: Hemisphere with no clear ocularity

**Figure S3.1:**
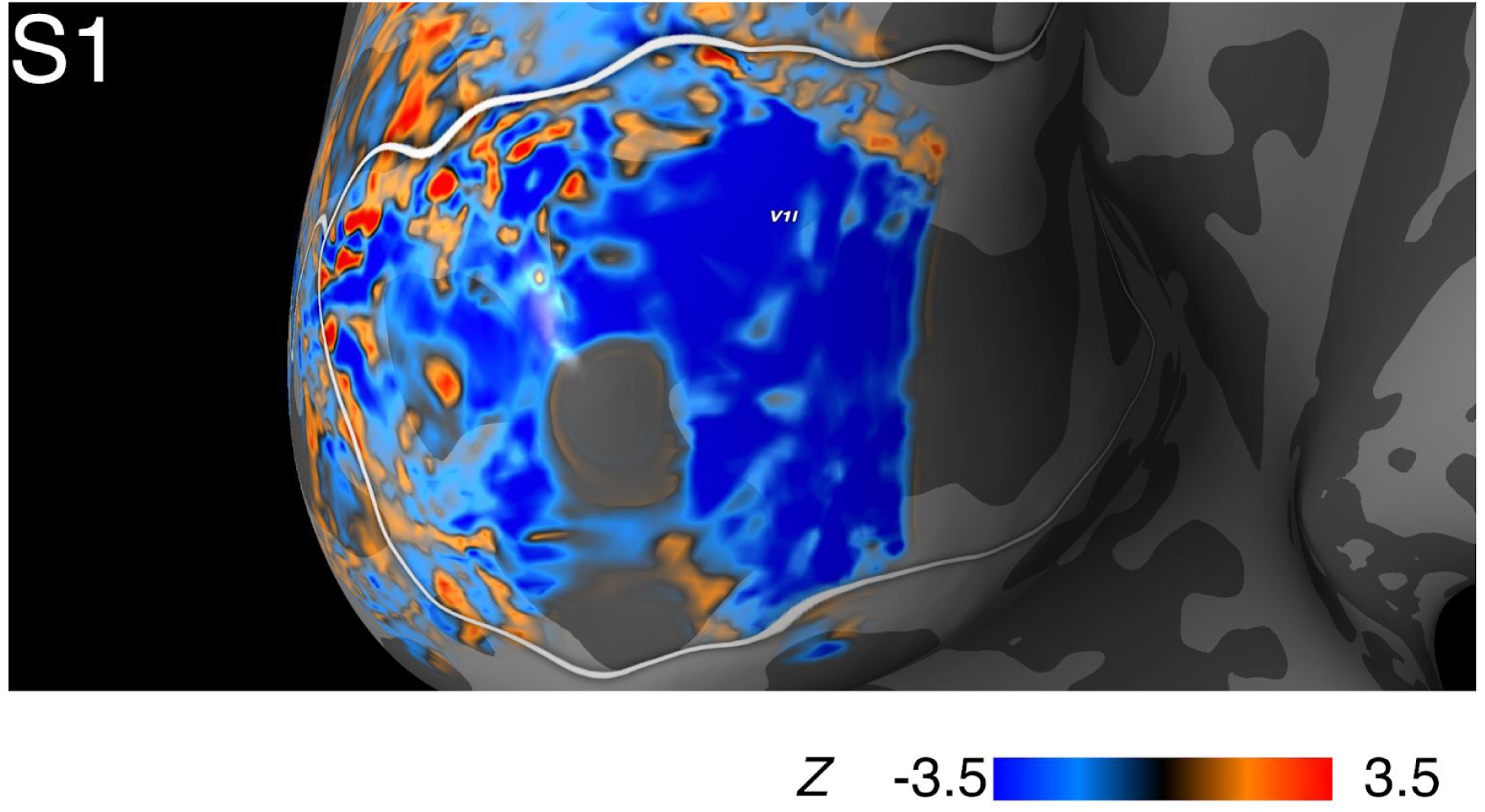
the left hemisphere of subject S1 shows a region of monocular representation. The “gray holes” without z-values lie more rostral than the surrounding parts of V1 and are therefore out of the field-of-view of the MRI protocol.

For one subject, S1, we found a large area of monocularity that does not adhere to the columnar pattern we would expect (see Figure S3.1). This monocular region was largely restricted more eccentric parts of the visual field. We do not know what caused this abnormality. One possibility is that this is an actual individual variation of the neural architecture. However, the subject did not report any problems with stereovision. Another possibility is that there was a retinotopic effect of the dichoptic stimulus presentation in the eccentric parts of the visual field. The “gray holes” without z-values lie more rostral than the other parts of V1 and are therefore out of the field-of-view of the receiver coil.

## Footnotes

1 https://github.com/VU-Cog-Sci/odc_mapping

2 https://openneuro.org/datasets/ds002295

3 https://github.com/kwagstyl/surface_tools

4 https://github.com/spinoza-centre/spynoza/blob/7t_hires/spynoza/hires/workflows.py

## References

Abadi M, Agarwal A, Barham P, Brevdo E, Chen Z, Citro C, Corrado GS, Davis A, Dean J, Devin M, Ghemawat S, Goodfellow I, Harp A, Irving G, Isard M, Jia Y, Jozefowicz R, Kaiser L, Kudlur M, Levenberg J, Mane D, Monga R, Moore S, Murray D, Olah C, Schuster M, Shlens J, Steiner B, Sutskever I, Talwar K, Tucker P, Vanhoucke V, Vasudevan V, Viegas F, Vinyals O, Warden P, Wattenberg M, Wicke M, Yu Y, Zheng X. 2016. TensorFlow: Large-Scale Machine Learning on Heterogeneous Distributed Systems. arXiv [csDC].

Adams DL, Horton JC. 2009. Ocular Dominance Columns: Enigmas and Challenges. The Neuroscientist. doi: 10.1177/1073858408327806

Adams DL, Sincich LC, Horton JC. 2007. Complete pattern of ocular dominance columns in human primary visual cortex. J Neurosci 27 :10391–10403.

Anderson JC, Martin KAC. 2009. The synaptic connections between cortical areas V1 and V2 in macaque monkey. J Neurosci 29 :11283–11293.

Andersson JLR, Skare S, Ashburner J. 2003. How to correct susceptibility distortions in spin-echo echo-planar images: application to diffusion tensor imaging. Neuroimage 20 :870–888.

Avants BB, Epstein CL, Grossman M, Gee JC. 2008. Symmetric diffeomorphic image registration with cross-correlation: evaluating automated labeling of elderly and neurodegenerative brain. Med Image Anal 12 :26–41.

Avants BB, Tustison NJ, Stauffer M, Song G, Wu B, Gee JC. 2014. The Insight ToolKit image registration framework. Front Neuroinform 8 :44.

Bartholomew-Biggs M, Brown S, Christianson B, Dixon L. 2000. Automatic differentiation of algorithms. J Comput Appl Math 124 :171–190.

Bastos AM, Usrey WM, Adams RA, Mangun GR, Fries P, Friston KJ. 2012. Canonical microcircuits for predictive coding. Neuron 76 :695–711.

Bazin P-L, Weiss M, Dinse J, Schäfer A, Trampel R, Turner R. 2014. A computational framework for ultra-high resolution cortical segmentation at 7Tesla. Neuroimage 93 Pt 2 :201–209.

Behzadi Y, Restom K, Liau J, Liu TT. 2007. A component based noise correction method (CompCor) for BOLD and perfusion based fMRI. Neuroimage 37 :90–101.

Bogovic JA, Prince JL, Bazin P-L. 2013. A Multiple Object Geometric Deformable Model for Image Segmentation. Comput Vis Image Underst 117 :145–157.

Brascamp J, Sterzer P, Blake R, Knapen T. 2018. Multistable Perception and the Role of the Frontoparietal Cortex in Perceptual Inference. Annu Rev Psychol 69 :77–103.

Caan MWA, Bazin P-L, Marques JP, de Hollander G, Dumoulin SO, van der Zwaag W. 2019. MP2RAGEME: T, T, and QSM mapping in one sequence at 7 tesla. Hum Brain Mapp 40 :1786–1798.

Chaimow D, Ugurbil K, Shmuel A. 2018a. Optimization of functional MRI for detection, decoding and high-resolution imaging of the response patterns of cortical columns. Neuroimage 164 :67–99.

Chaimow D, Yacoub E, Uğurbil K, Shmuel A. 2018b. Spatial specificity of the functional MRI blood oxygenation response relative to neuronal activity. Neuroimage 164:32–47.

Chaimow D, Yacoub E, Ugurbil K, Shmuel A. 2011. Modeling and analysis of mechanisms underlying fMRI-based decoding of information conveyed in cortical columns. Neuroimage 56 :627–642.

Cheng K, Waggoner RA, Tanaka K. 2001. Human ocular dominance columns as revealed by high-field functional magnetic resonance imaging. Neuron 32 :359–374.

Constantinople CM, Bruno RM. 2013. Deep cortical layers are activated directly by thalamus. Science 340 :1591–1594.

Corbitt PT, Ulloa A, Horwitz B. 2018. Simulating laminar neuroimaging data for a visual delayed match-to-sample task. NeuroImage. doi: 10.1016/j.neuroimage.2018.02.037

Corliss G, Faure C, Griewank A, Hascoet L, Naumann U. 2002. Automatic Differentiation of Algorithms: From Simulation to Optimization. Springer Science & Business Media.

Cox RW. 1996. AFNI: software for analysis and visualization of functional magnetic resonance neuroimages. Comput Biomed Res 29 :162–173.

Crandall SR, Patrick SL, Cruikshank SJ, Connors BW. 2017. Infrabarrels Are Layer 6 Circuit Modules in the Barrel Cortex that Link Long-Range Inputs and Outputs. Cell Rep 21 :3065–3078.

Dale AM, Fischl B, Sereno MI. 1999. Cortical surface-based analysis. I. Segmentation and surface reconstruction. Neuroimage 9 :179–194.

Dechent P, Frahm J. 2000. Direct mapping of ocular dominance columns in human primary visual cortex. Neuroreport 11 :3247–3249.

de Hollander G. 2018. Pymp2rage. GitHub. doi: 10.5281/zenodo.1476977

De Martino F, Moerel M, Ugurbil K, Goebel R, Yacoub E, Formisano E. 2015. Frequency preference and attention effects across cortical depths in the human primary auditory cortex. Proc Natl Acad Sci U S A 112 :16036–16041.

De Martino F, Yacoub E, Kemper V, Moerel M, Uludağ K, De Weerd P, Ugurbil K, Goebel R, Formisano E. 2018. The impact of ultra-high field MRI on cognitive and computational neuroimaging. Neuroimage 168 :366–382.

Denfield GH, Ecker AS, Shinn TJ, Bethge M, Tolias AS. 2018. Attentional fluctuations induce shared variability in macaque primary visual cortex. Nat Commun 9 :2654.

DeSimone K, Rokem A, Schneider K. 2016. popeye: a population receptive field estimation tool. The Journal of Open Source Software. doi: 10.21105/joss.00103

Dougherty K, Cox MA, Westerberg JA, Maier A. 2019. Binocular Modulation of Monocular V1 Neurons. Curr Biol 29 :381–391.e4.

Dumoulin SO, Fracasso A, van der Zwaag W, Siero JCW, Petridou N. 2018. Ultra-high field MRI: Advancing systems neuroscience towards mesoscopic human brain function. Neuroimage 168 :345–357.

Dumoulin SO, Wandell BA. 2008. Population receptive field estimates in human visual cortex. Neuroimage 39 :647–660.

Esteban O, Markiewicz CJ, Blair RW, Moodie CA, Isik AI, Erramuzpe A, Kent JD, Goncalves M, DuPre E, Snyder M, Oya H, Ghosh SS, Wright J, Durnez J, Poldrack RA, Gorgolewski KJ. 2019. fMRIPrep: a robust preprocessing pipeline for functional MRI. Nat Methods 16 :111–116.

Fischl B, Sereno MI, Dale AM. 1999. Cortical surface-based analysis. II: Inflation, flattening, and a surface-based coordinate system. Neuroimage 9 :195–207.

Fonov VS, Evans AC, McKinstry RC, Almli CR, Collins DL. 2009. Unbiased nonlinear average age-appropriate brain templates from birth to adulthood. NeuroImage. doi: 10.1016/s1053-8119(09)70884-5

Forsyth DA, Ponce J. 2015. Computer Vision: A Modern Approach: A Modern Approach. Pearson Higher Ed.

Fracasso A, Petridou N, Dumoulin SO. 2016. Systematic variation of population receptive field properties across cortical depth in human visual cortex. Neuroimage 139 :427–438.

Friston KJ, Williams S, Howard R, Frackowiak RS, Turner R. 1996. Movement-related effects in fMRI time-series. Magn Reson Med 35 :346–355.

Gao JS, Huth AG, Lescroart MD, Gallant JL. 2015. Pycortex: an interactive surface visualizer for fMRI. Front Neuroinform 9 :23.

Gelman A, Hwang J, Vehtari A. 2014. Understanding predictive information criteria for Bayesian models. Stat Comput 24:997–1016.

Goncalves NR, Welchman AE. 2017. “What Not” Detectors Help the Brain See in Depth. Curr Biol 27 :1403–1412.e8.

Goodyear BG, Menon RS. 2001. Brief visual stimulation allows mapping of ocular dominance in visual cortex using fMRI. Hum Brain Mapp 14 :210–217.

Gorgolewski K, Burns CD, Madison C, Clark D, Halchenko YO, Waskom ML, Ghosh SS. 2011. Nipype: a flexible, lightweight and extensible neuroimaging data processing framework in python. Front Neuroinform 5 :13.

Greve DN, Fischl B. 2009. Accurate and robust brain image alignment using boundary-based registration. Neuroimage 48 :63–72.

Han X, Pham DL, Tosun D, Rettmann ME, Xu C, Prince JL. 2004. CRUISE: Cortical reconstruction using implicit surface evolution. NeuroImage. doi: 10.1016/j.neuroimage.2004.06.043

Havlicek M, Uludağ K. 2020. A dynamical model of the laminar BOLD response. Neuroimage 204 :116209.

Haxby JV, Gobbini MI, Furey ML, Ishai A, Schouten JL, Pietrini P. 2001. Distributed and overlapping representations of faces and objects in ventral temporal cortex. Science 293 :2425–2430.

Haynes J-D, Deichmann R, Rees G. 2005. Eye-specific effects of binocular rivalry in the human lateral geniculate nucleus. Nature 438 :496–499.

Haynes J-D, Rees G. 2006. Neuroimaging: decoding mental states from brain activity in humans. Nat Rev Neurosci 7 :523.

Haynes J-D, Rees G. 2005. Predicting the stream of consciousness from activity in human visual cortex. Curr Biol 15 :1301–1307.

Heinzle J, Koopmans PJ, den Ouden HEM, Raman S, Stephan KE. 2016. A hemodynamic model for layered BOLD signals. Neuroimage 125 :556–570.

Hoerl AE, Kennard RW. 1970. Ridge Regression: Applications to Nonorthogonal Problems. Technometrics. doi: 10.2307/1267352

Hoffman MD, Gelman A. 2014. The No-U-Turn sampler: adaptively setting path lengths in Hamiltonian Monte Carlo. J Mach Learn Res 15 :1593–1623.

Hubel DH, Wiesel TN. 1969. Anatomical demonstration of columns in the monkey striate cortex. Nature 221 :747–750.

Huntenburg JM, Steele CJ, Bazin P-L. 2018. Nighres: processing tools for high-resolution neuroimaging. Gigascience 7. doi: 10.1093/gigascience/giy082

Jenkinson M. 2002. Improved Optimization for the Robust and Accurate Linear Registration and Motion Correction of Brain Images. NeuroImage. doi: 10.1016/s1053-8119(02)91132-8

Jenkinson M, Bannister P, Brady M, Smith S. 2002. Improved Optimization for the Robust and Accurate Linear Registration and Motion Correction of Brain Images. NeuroImage. doi: 10.1006/nimg.2002.1132

Jenkinson M, Beckmann CF, Behrens TEJ, Woolrich MW, Smith SM. 2012. FSL. NeuroImage. doi: 10.1016/j.neuroimage.2011.09.015

Jenkinson, M., Pechaud, M., & Smith, S. 2005. BET2: MR-based estimation of brain, skull and scalp surfacesEleventh Annual Meeting of the Organization for Human Brain Mapping. doi: 10.1016/s1053-8119(08)70003-x

Kamitani Y, Tong F. 2005. Decoding the visual and subjective contents of the human brain. Nat Neurosci 8 :679–685.

Kay K, Jamison KW, Vizioli L, Zhang R, Margalit E, Ugurbil K. 2019. A critical assessment of data quality and venous effects in sub-millimeter fMRI. Neuroimage 189 :847–869.

Klein A, Ghosh SS, Bao FS, Giard J, Häme Y, Stavsky E, Lee N, Rossa B, Reuter M, Chaibub Neto E, Keshavan A. 2017. Mindboggling morphometry of human brains. PLoS Comput Biol 13 :e1005350.

Koopmans PJ, Barth M, Norris DG. 2010. Layer-specific BOLD activation in human V1. Hum Brain Mapp 31 :1297–1304.

Kruschke J. 2014. Doing Bayesian Data Analysis: A Tutorial with R, JAGS, and Stan. Academic Press.

Kuehn E, Sereno MI. 2018. Modelling the Human Cortex in Three Dimensions. Trends Cogn Sci 22 :1073–1075.

Lawrence SJ, Norris DG, de Lange FP. 2019. Dissociable laminar profiles of concurrent bottom-up and top-down modulation in the human visual cortex. Elife 8. doi: 10.7554/eLife.44422

Ledoit O, Wolf M. 2003. Improved estimation of the covariance matrix of stock returns with an application to portfolio selection. Journal of Empirical Finance. doi: 10.1016/s0927-5398(03)00007-0

Leopold DA, Logothetis NK. 1996. Activity changes in early visual cortex reflect monkeys’ percepts during binocular rivalry. Nature 379 :549–553.

Levelt WJM. 1965. On binocular rivalry.

Markuerkiaga I, Barth M, Norris DG. 2016. A cortical vascular model for examining the specificity of the laminar BOLD signal. Neuroimage 132 :491–498.

Marquardt I, Schneider M, Gulban OF, Ivanov D, Uludağ K. 2018. Cortical depth profiles of luminance contrast responses in human V1 and V2 using 7 T fMRI. Human Brain Mapping. doi: 10.1002/hbm.24042

Marques JP, Gruetter R. 2013. New developments and applications of the MP2RAGE sequence--focusing the contrast and high spatial resolution R1 mapping. PLoS One 8 :e69294.

Marques JP, Kober T, Krueger G, van der Zwaag W, Van de Moortele P-F, Gruetter R. 2010. MP2RAGE, a self bias-field corrected sequence for improved segmentation and T1-mapping at high field. Neuroimage 49 :1271–1281.

Marques JP, Norris DG. 2018. How to choose the right MR sequence for your research question at 7 T and above? Neuroimage 168 :119–140.

McCarthy P. 2019. FSLeyes. doi: 10.5281/zenodo.3403671

Menon RS, Ogawa S, Strupp JP, Uğurbil K. 1997. Ocular dominance in human V1 demonstrated by functional magnetic resonance imaging. J Neurophysiol 77 :2780–2787.

Merola A, Weiskopf N. 2018. Modelling the laminar GE-BOLD signal: integrating anatomical, physiological and methodological factorsJoint Annual Meeting ISMRM-ESMRMB,.

Moon C-H, Fukuda M, Park S-H, Kim S-G. 2007. Neural interpretation of blood oxygenation level-dependent fMRI maps at submillimeter columnar resolution. J Neurosci 27 :6892–6902.

Muckli L, De Martino F, Vizioli L, Petro LS, Smith FW, Ugurbil K, Goebel R, Yacoub E. 2015. Contextual Feedback to Superficial Layers of V1. Curr Biol 25 :2690–2695.

Mur M, Bandettini PA, Kriegeskorte N. 2009. Revealing representational content with pattern-information fMRI--an introductory guide. Soc Cogn Affect Neurosci 4 :101–109.

Nistats. 2019. https://nistats.github.io/index.html

Norman KA, Polyn SM, Detre GJ, Haxby JV. 2006. Beyond mind-reading: multi-voxel pattern analysis of fMRI data. Trends Cogn Sci 10 :424–430.

Nunez-Elizalde AO, Huth AG, Gallant JL. 2019. Voxelwise encoding models with non-spherical multivariate normal priors. Neuroimage 197 :482–492.

O’Connor DH, Fukui MM, Pinsk MA, Kastner S. 2002. Attention modulates responses in the human lateral geniculate nucleus. Nat Neurosci 5 :1203–1209.

O’Toole AJ, Jiang F, Abdi H, Pénard N, Dunlop JP, Parent MA. 2007. Theoretical, statistical, and practical perspectives on pattern-based classification approaches to the analysis of functional neuroimaging data. J Cogn Neurosci 19 :1735–1752.

Petridou N, Italiaander M, van de Bank BL, Siero JCW, Luijten PR, Klomp DWJ. 2013. Pushing the limits of high-resolution functional MRI using a simple high-density multi-element coil design. NMR Biomed 26 :65–73.

Petro LS, Vizioli L, Muckli L. 2014. Contributions of cortical feedback to sensory processing in primary visual cortex. Front Psychol 5 :1223.

Poldrack RA, Gorgolewski KJ, Varoquaux G. 2019. Computational and Informatic Advances for Reproducible Data Analysis in Neuroimaging. Annu Rev Biomed Data Sci 2 :119–138.

Polimeni JR, Renvall V, Zaretskaya N, Fischl B. 2018. Analysis strategies for high-resolution UHF-fMRI data. Neuroimage 168 :296–320.

Power JD, Barnes KA, Snyder AZ, Schlaggar BL, Petersen SE. 2012. Spurious but systematic correlations in functional connectivity MRI networks arise from subject motion. Neuroimage 59 :2142–2154.

Power JD, Mitra A, Laumann TO, Snyder AZ, Schlaggar BL, Petersen SE. 2014. Methods to detect, characterize, and remove motion artifact in resting state fMRI. Neuroimage 84 :320–341.

Puckett AM, Aquino KM, Robinson PA, Breakspear M, Schira MM. 2016. The spatiotemporal hemodynamic response function for depth-dependent functional imaging of human cortex. Neuroimage 139 :240–248.

Rockland KS, Pandya DN. 1979. Laminar origins and terminations of cortical connections of the occipital lobe in the rhesus monkey. Brain Res 179 :3–20.

Rockland KS, Virga A. 1989. Terminal arbors of individual “feedback” axons projecting from area V2 to V1 in the macaque monkey: a study using immunohistochemistry of anterogradely transported Phaseolus vulgaris-leucoagglutinin. J Comp Neurol 285 :54–72.

Salvatier J, Wiecki TV, Fonnesbeck C. 2016. Probabilistic programming in Python using PyMC3. PeerJ Comput Sci 2 :e55.

Savitzky A, Golay MJE. 1964. Smoothing and Differentiation of Data by Simplified Least Squares Procedures. Analytical Chemistry. doi: 10.1021/ac60214a047

Schneider M, Kemper VG, Emmerling TC, De Martino F, Goebel R. 2019. Columnar clusters in the human motion complex reflect consciously perceived motion axis. Proc Natl Acad Sci U S A 116 :5096–5101.

Schurger A. 2009. A very inexpensive MRI-compatible method for dichoptic visual stimulation. J Neurosci Methods 177 :199–202.

Self MW, van Kerkoerle T, Goebel R, Roelfsema PR. 2019. Benchmarking laminar fMRI: Neuronal spiking and synaptic activity during top-down and bottom-up processing in the different layers of cortex. Neuroimage 197 :806–817.

Shmuel A, Yacoub E, Chaimow D, Logothetis NK, Ugurbil K. 2007. Spatio-temporal point-spread function of fMRI signal in human gray matter at 7 Tesla. NeuroImage. doi: 10.1016/j.neuroimage.2006.12.030

Siero JCW, Petridou N, Hoogduin H, Luijten PR, Ramsey NF. 2011. Cortical depth-dependent temporal dynamics of the BOLD response in the human brain. J Cereb Blood Flow Metab 31 :1999–2008.

Smyser CD, Inder TE, Shimony JS, Hill JE, Degnan AJ, Snyder AZ, Neil JJ. 2010. Longitudinal analysis of neural network development in preterm infants. Cereb Cortex 20 :2852–2862.

Stephan KE, Petzschner FH, Kasper L, Bayer J, Wellstein KV, Stefanics G, Pruessmann KP, Heinzle J. 2019. Laminar fMRI and computational theories of brain function. Neuroimage 197 :699–706.

Tong F, Meng M, Blake R. 2006. Neural bases of binocular rivalry. Trends Cogn Sci 10 :502–511.

Tong F, Pratte MS. 2012. Decoding patterns of human brain activity. Annu Rev Psychol 63 :483–509.

Tootell RB, Hamilton SL, Silverman MS, Switkes E. 1988. Functional anatomy of macaque striate cortex. I. Ocular dominance, binocular interactions, and baseline conditions. J Neurosci 8 :1500–1530.

Trattnig S, Springer E, Bogner W, Hangel G, Strasser B, Dymerska B, Cardoso PL, Robinson SD. 2018. Key clinical benefits of neuroimaging at 7T. Neuroimage 168 :477–489.

Tsuchiya N, Koch C. 2005. Continuous flash suppression reduces negative afterimages. Nat Neurosci 8 :1096–1101.

Tustison NJ, Avants BB, Cook PA, Zheng Y, Egan A, Yushkevich PA, Gee JC. 2010. N4ITK: improved N3 bias correction. IEEE Trans Med Imaging 29 :1310–1320.

van Bergen RS, Jehee JFM. 2019. Improved methods for decoding sensory uncertainty from activity in the human visual cortexVSS Conference 2019. Presented at the VSS 2019.

van Bergen RS, Ma WJ, Pratte MS, Jehee JFM. 2015. Sensory uncertainty decoded from visual cortex predicts behavior. Nature Neuroscience. doi: 10.1038/nn.4150

van der Zwaag W, Schäfer A, Marques JP, Turner R, Trampel R. 2016. Recent applications of UHF-MRI in the study of human brain function and structure: a review. NMR Biomed 29 :1274–1288.

van Kerkoerle T, Self MW, Roelfsema PR. 2017. Layer-specificity in the effects of attention and working memory on activity in primary visual cortex. Nat Commun 8 :13804.

Waehnert MD, Dinse J, Weiss M, Streicher MN, Waehnert P, Geyer S, Turner R, Bazin P-L. 2014. Anatomically motivated modeling of cortical laminae. Neuroimage 93 Pt 2 :210–220.

Wandell BA, Winawer J. 2011. Imaging retinotopic maps in the human brain. Vision Research. doi: 10.1016/j.visres.2010.08.004

Watanabe S. 2013. A Widely Applicable Bayesian Information Criterion. J Mach Learn Res 14 :867–897.

Wheatstone C. 1838. Contributions to the physiology of vision. —Part the first. On some remarkable, and hitherto unobserved, phenomena of binocular vision. Philosophical Transactions of the Royal Society of London 128 :371–394.

Wunderlich K, Schneider KA, Kastner S. 2005. Neural correlates of binocular rivalry in the human lateral geniculate nucleus. Nat Neurosci 8 :1595–1602.

Yacoub E, Shmuel A, Logothetis N, Uğurbil K. 2007. Robust detection of ocular dominance columns in humans using Hahn Spin Echo BOLD functional MRI at 7 Tesla. Neuroimage 37 :1161–1177.

Zhang N, Zhu X-H, Zhang Y, Park J-K, Chen W. 2010. High-resolution fMRI mapping of ocular dominance layers in cat lateral geniculate nucleus. Neuroimage 50 :1456–1463.

Zhang P, Jamison K, Engel S, He B, He S. 2011. Binocular rivalry requires visual attention. Neuron 71 :362–369.

Zhang P, Jiang Y, He S. 2012. Voluntary attention modulates processing of eye-specific visual information. Psychol Sci 23 :254–260.

Zhang Y, Brady M, Smith S. 2001. Segmentation of brain MR images through a hidden Markov random field model and the expectation-maximization algorithm. IEEE Trans Med Imaging 20 :45–57.

